# Mechanical model of muscle contraction 2. Kinematic and dynamic aspects of a myosin II head during the working stroke

**DOI:** 10.1101/2019.12.16.878801

**Authors:** S. Louvet

## Abstract

The condition of a myosin II head during which force and movement are generated is commonly referred to as Working Stroke (WS). During the WS, the myosin head is mechanically modelled by 3 two by two articulated segments, the motor domain (S1a) strongly fixed to an actin molecule, the lever (S1b) on which a motor moment is exerted, and the rod (S2) pulling the myosin filament (Mfil). When the half-sarcomere (hs) is shortened or lengthened by a few nanometers, it is assumed that the lever of a myosin head in WS state moves in a fixed plane including the longitudinal axis of the actin filament (Afil). As a result, the 5 rigid segments, i.e. Afil, S1a, S1b, S2 and Mfil, follow deterministic and configurable trajectories. The orientation of S1b in the fixed plane is characterized by the angle θ. After deriving the geometric equations singularizing the WS state, we obtain an analytical relationship between the hs shortening velocity (u) and the angular velocity of the lever 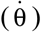. The principles of classical mechanics applied to the 3 solids, S1a, S1b and S2, lead to a relationship between the motor moment exerted on the lever (M_B_) and the tangential force dragging the actin filament (T_A_). We distinguish θ_up_ and θ_down_, the two boundaries framing the angle θ during the WS, relating to *up* and *down* conformations. With the usual data assigned to the cross-bridge elements, a linearization procedure of the relationships between u and 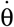, on the one hand, and between M_B_ and T_A_, on the other hand, is performed. This algorithmic optimization leads to theoretical values of θ_up_ and θ_down_ equal to +28° (−28°) and −42° (+42°) respectively with a variability of ±5° in a hs on the right (left), data in accordance with the commonly accepted experimental values for vertebrate muscle fibers.

## Introduction

The mechanical study of human movement requires modelling the body as a deformable material set composed of articulated rigid segments. In this context, the calculation of the powers of the internal actions, i.e. muscular, is similar to that of the powers of the articular moments where only the rotational movements are important. For example, to simulate the walking movement, rotational motors are positioned at the joints of the members of a humanoid robot and the combination of their actions ultimately reproduce a translation movement of the robot’s centre of gravity. In 1993, I Rayment and his co-authors [1,2] established that the dynamics of a myosin head is associated with the rotational movement of lever S1b, indicating that the linear shortening of a hs results from the collective rotational movements of the levers. The analogy present in the phrases “*lever arm theory*” or “*model of swinging lever arm*” suggests studying muscle fiber as a deformable material set to which the general principles of classical mechanics are applied.

Our model assumes that the movement of lever S1b belonging to a WS myosin head is carried out in a fixed plane during the entire WS state. This fourth hypothesis leads to the emblematic values of θ_up_ and θ_down_ and justifies the impossibility given to the two heads of a myosin molecule to be simultaneously in WS state. The hypothesis n° 4 is demonstrated in paragraph F.4 of Supplement S2.F because it proves to be the necessary and sufficient condition for the the linear moment principle and the work-energy theorem applied to the muscle fiber to provide identical formulations, a key tenet of classical mechanics. This paper is the first step that leads to the calculation of the tension after isometric tetanization and after phase 1 of a length step (Paper 4), to the expression of the tension increase during the last 3 phases of a length step (Paper 5), to the determination of the tension after a succession of length steps (Paper 6) and finally to the Force/Velocity relationship examined in Paper 1.

## Methods

When the myosin head is strongly bound to actin, the 5 segments, Afil, S1a, S1b, S2 and Mfil, form a poly-articulated chain usually called cross-bridge (Fig 1a). The 4 joints of the chain are represented by the 4 points A, B, C and D (Fig 1d). The O_Afil_X axis is the longitudinal axis of the actin filament and the O_Afil_Y° axis is the axis perpendicular to O_Afil_X passing through the centers of the adjacent actin and myosin filaments (Fig 1b).

**Fig 1.**
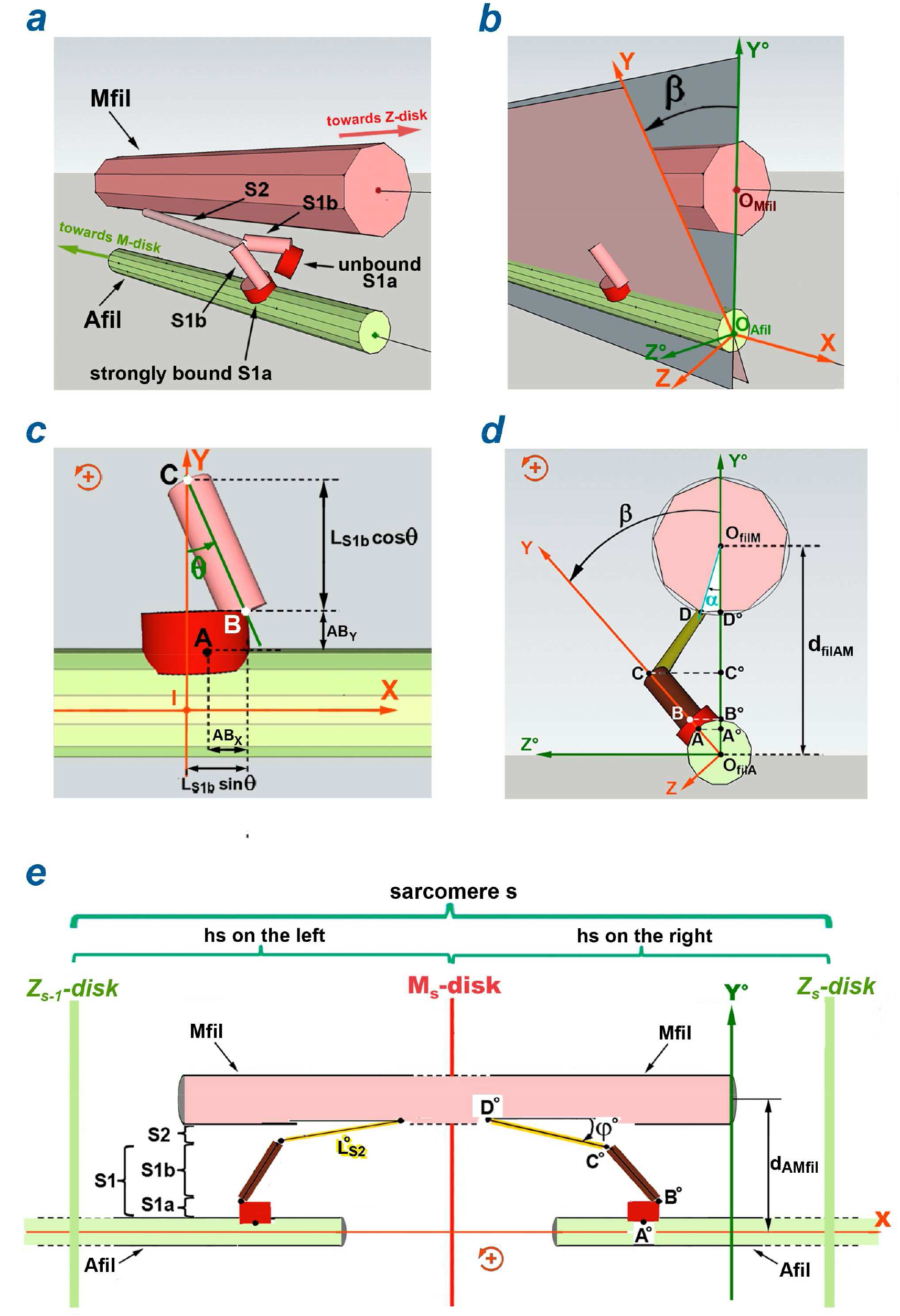
Geometric characteristics of a WS myosin head. (a) Myosin molecule, one of whose 2 heads is in WS forming a cross-bridge between the myosin filament (Mfil) and the actin filament (Afil). (b) Definition of the angle β in the O_Afil_Y°Z° plane. (c) Definition of the θ angle in the IXY plane where point I is the projection of C on O_Afil_X according to O_Afil_Y. (d) Cross-section in the O_Afil_Y°Z° plane where A°, B°, C° and D° are the orthogonal projections of A, B, C and D in the O_Afil_XY° plane. (e) Orthogonal projection of a schematized sarcomere in the O_Afil_XY° plane.

Classically the WS state presented by I. Rayment [1] is based on 3 conditions: (1) the segments S1a, S1b and S2 are rigid, (2) S1a is strongly or “stereo-specifically” attached to an actin molecule of Afil, (3) a motor-moment is exerted on S1b and consequently a tensile force is applied to the Mfil via S2.

We propose a fourth condition which assumes that during the WS, the lever S1b represented by the segment BC moves in the fixed plane O_Afil_XY forming the angle β with the plane O_Afil_XY (Figs 1b and 1d), the orientation of the lever S1b in the plane O_Afil_XY being characterized by the angle θ (Fig 1c).

The four supplements associated with the article and cited in the text are entitled S2.C, S2.D, S2.E and S2.F.

### Kinematics of a myosin head in WS

Our assumption allows the calculation of the positions of the 3 segments S1a, S1b and S2 (see paragraph D.1 of Supplement S2.D). By temporal derivation, we obtain a relationship between the relative hs shortening velocity (u) and the angular velocity of S1b 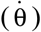:

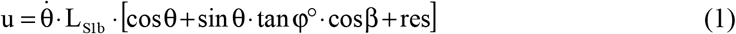

where 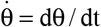; L_S1b_ is the S1b length with L_S1b_=|BC|; φ° is the instantaneous angle between D°Y and D°C° in the O_Mfil_XY° plane (Fig 1e), points C° and D° are the orthogonal projections of C and D on the O_Afil_XY° plane (Fig 1d); res is a “residual” function taking instantaneous values close to 0, whose formula is explained in (D10) in the Supplement S2.D.

Since φ° depends on θ, α and β according to the equality (D3) of Supplement S2.D, we introduce the analytical function:

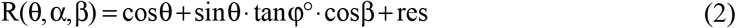

With (2), equality (1) is rewritten:

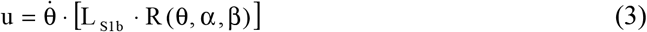

### Dynamics of a WS head located in a hs that shortens at constant speed

Our model is applied to a hs of the muscle fiber in isometric conditions or shortening at steady slow speed. In this case, it is attested with the calculations developed in Supplement S1.C that the only mechanical actions present in the hs are the linking forces and moments of the poly-articulated chains (Afil + S1a + S1b + S2 + Mfil) where S1a, S1b and S2 belong to the heads in WS state (Fig 2). A motor-moment (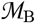) is enforced to point B symbolizing the pivot joint between S1a and S1b during the WS. The moments in C and D are zero.

**Fig 2.**
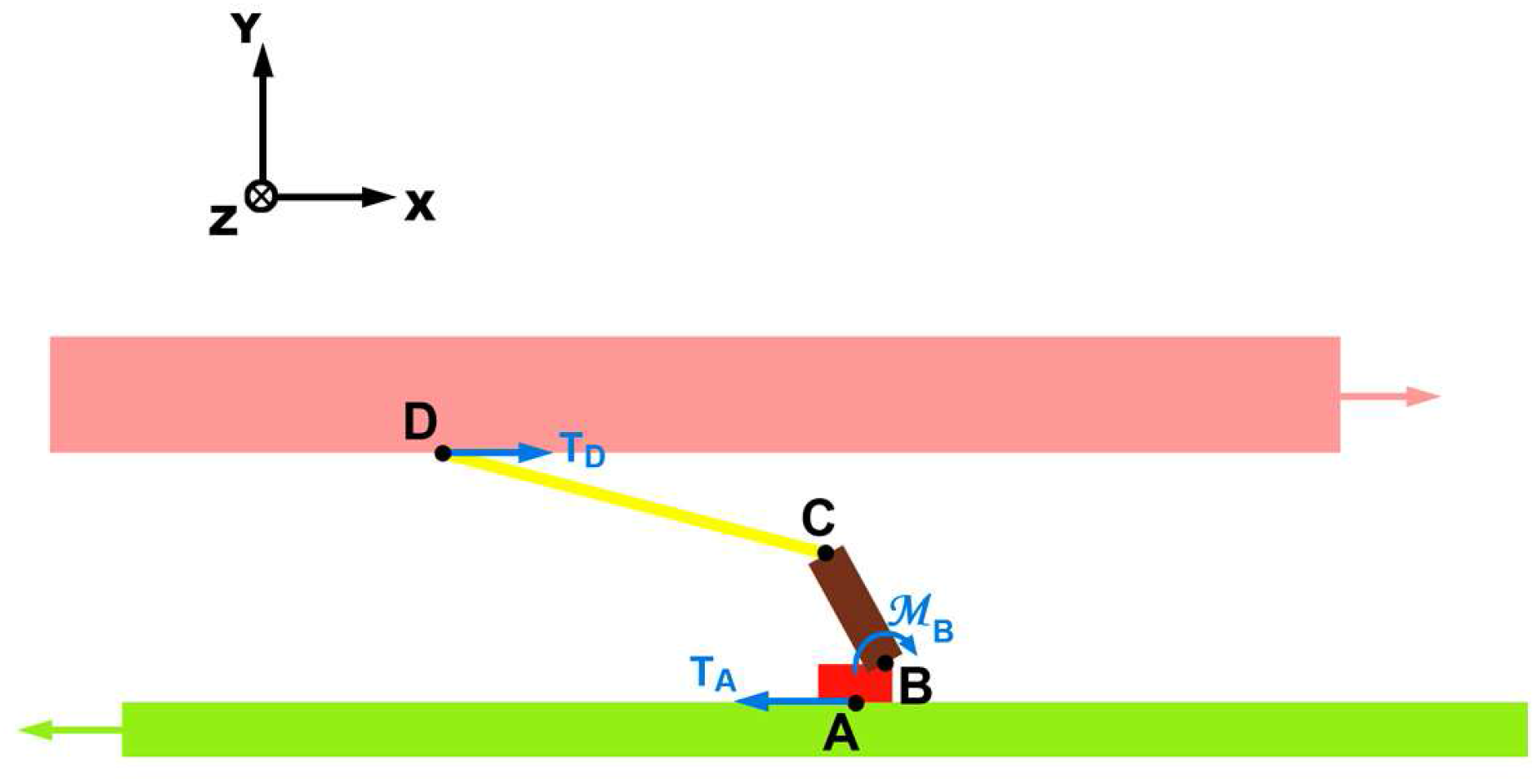
Actions of the inter-segmentary links of a myosin head in Working Stroke.

Among the forces exerted on S1a, S1b and S2, only the tangential components T_A_ and T_D_ relating to points A and D and the 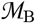 moment exerted at point B are shown in Fig 2; see Fig D2 of Supplement S2.D for a complete representation of the actions involved.

The linear and angular moment principles are implemented to a WS myosin head located in a hs on the right; see paragraph D.2 of Supplement S2.D. They provide the following relationships:

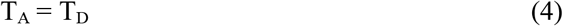

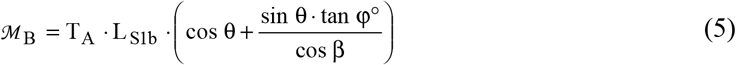

It is posed:

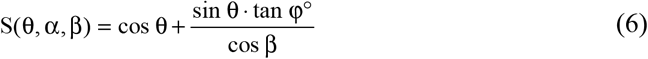

With (6), equality (5) is reformulated:

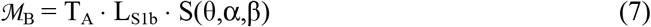

### Calculation of θ_up_

The purpose of this work is to specify the values of θ_down_ and θ_up_ which optimize the linearization of the relationship (3), which is equivalent to seeking the constancy of the function R(θ,α,β) between these 2 limits. The maximum variation of the θ angle during the WS (δθ_Max_) is given classically equal to 70° [2,3,4,5,6]. This value is recorded in Table 1 and only θ_down_ remains to be determined because:

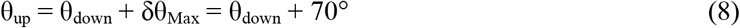

**Table 1.** Geometric characteristics of a myosin II head in WS in a vertebrate half-sarcomere on the right.

| Acronym | Definition | Value | Variability range |
| --- | --- | --- | --- |
| $AB_X$ | Distance between A and B according to OX | 2 nm | [0 nm ; 5 nm] |
| $AB_Y$ | Distance between A and B according to OY | 1.5 nm | [0.5 nm ; 3 nm] |
| $L_{S1b}$ | Length of the lever S1b | 10 nm | [8 nm; 11 nm] |
| $L_{S2}$ | Length of the rod S2 | 50 nm | [30 nm ; 65 nm] |
| $d_{2Mfil}^{(1)}$ | Distance between the centers of 2 adjacent Mfil | 46 nm | [40 nm ; 55 nm] |
| $d_{AMfi}^{(1)}$ | Distance between the centers of adjacent Afil and Mfil | 26.6 nm | [23 nm ; 32 nm] |
| $\delta X_{Max}$ | Stroke size or maximum step during the WS | 11.5 nm | [10 nm; 12 nm] |
| $\delta \theta_{Max}^{(2)}$ | Maximum variation of $\theta$ during the WS | 70° | [60° ; 80°] |

### Determination of R_0_(θ) and S_0_(θ)

When both α and β angles are zero, functions R(θ,0,0) and S(θ,0,0) are noted as R_0_(θ) and S_0_(θ), respectively. With the formula given in (D10) in Supplement S2.D, we note that the residual function (res) present in (1) and (2) is cancelled if β=0. Under these conditions, according to equations (2) and (6), the following relations are verified:

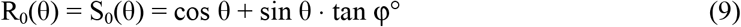

### Algorithmic methods

The algorithms work as follows: the θ_down_ angle is incremented from −90° to 0° with a pitch of 0.5°, i.e. a total of 181 θ_down_ values tested. At each pitch, the angle θ_up_ associated with θ_down_ is determined using equality (8).

1. **Minimization of deviations** At each iteration of θ_down_ the variable R_0_ is evaluated according to (9) for θ varying between θ_down_ and θ_up_ with a pitch of 0.5°. Then the average (R_WS_) of these 141 values of R_0_ is calculated. Finally, the 141 deviations between R_WS_ and the 141 values of R_0_ are characterized and summed. The value of θ_down_ for which the sum of differences is minimal is chosen.
2. **Zero slope** At each iteration of θ_down_ the slope of the regression line between the 2 variables R_0_ and θ is calculated, θ being incremented between θ_down_ and θ_up_ with a pitch of 0.5°. The program searches for the value of θ_down_ for which the slope cancels out or is closest to zero, i.e. the value for which R_0_ becomes independent of θ.

With respect to both methods, all calculations are performed using reference data [7.8] that are reported in the “Value” column of Table 1. In both cases, it must be ensured a posteriori that the calculation carried out between the limits θ_down_ and θ_up_ provides the value planned for δX_Max_ defined as following:

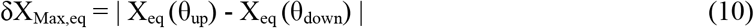

where X_eq_ is the exact calculation of the abscissa X according to equation (D5) of Supplement S2.D, equation reproduced below :

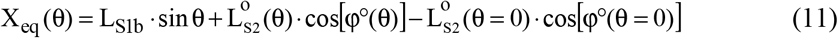

where the different terms present in the right member of (11) are explained in Supplement S2.D.

All calculation algorithms have been developed with Visual Basic 6 software.

### Statistics

Application of the linear regression is described in Methods section of Paper 1.

## Results

### Determination of θ_down_

The plot of the R_0_ function defined in (9) appears with a red line on Fig 3a. The deviation minimization algorithm applied to R_0_ described in the Methods section provides the value of θ_down_ in a hs on the right:

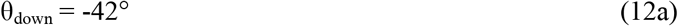

**Fig 3.**
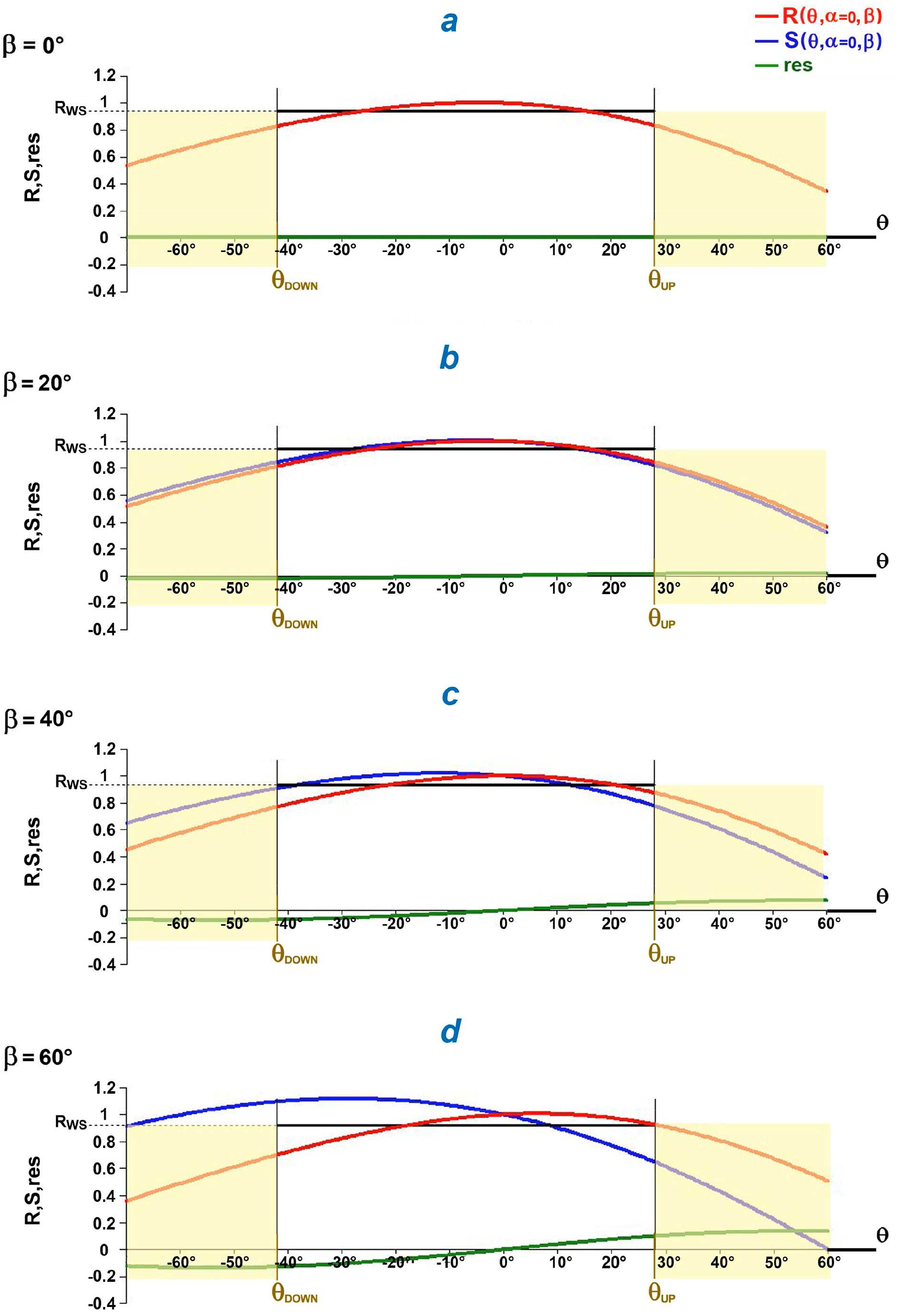
Evolutions of R(θ,α,β), S(θ,α,β) and res(θ,α,β) in function of θ varying between −70° and +60° according to 4 values from β with α=0°. (a) β = 0°; (b) β = 20°; (c) β = 40°; (d) β = 60°.

The *down* position of lever S1b at the end of the WS is also called as post-powerstroke, prerecovery, like-rigor or M*. Its angular value is given equal to 45° in modulus [9,10,11,12,13,14], which is close to the value found in (12a).

The *up* position of lever S1b at the beginning of the WS, also called as pre-powerstroke, postrecovery or M**, takes for orientation according to the relations (8) and (12a):

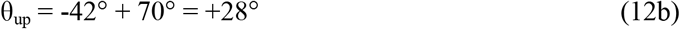

In a half-sarcomere on the left, the boundaries θ_down_ and θ_up_ are identical in module and opposite as a sign.

### Linearization of equations between θ_down_ and θ_up_

The averages of the two functions R_0_ and S_0_ between the two limits θ_down_ and θ_up_ determined by (12a) and (12b) are called R_WS_ and S_WS_, respectively. With the equality (9), the algorithmic calculations of R_WS_ and S_WS_ give (line β=0° of Table E1 of Supplement S2.E):

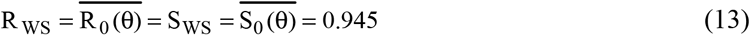

Between θ_down_ and θ_up_, equations (3) and (7) are approximated by the 2 respective constants R_WS_ and S_WS_, (Figs 3a, 3b, 3c and 3d; thick black line) such that:

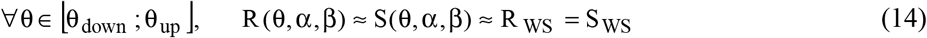

Using the equalities presented in (14), we proceed between θ_down_ and θ_up_ to the linearization of equations (3) and (7), respectively:

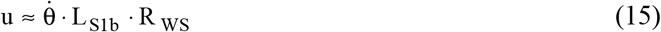

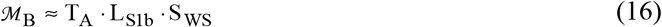

### Scope of validity of the linearization according to β

The angle β varies between −60° and +60°; see expression (G9) of Supplement S3.G of accompanying Paper 3. In equations (2), (3), (6), (7) and (9), only the cosine of β is involved, so the study of the positive values of β is sufficient. The functions R(θ,0,β), R(θ,0,β) and res(θ,0,β) are represented as a function of θ by red, blue and green lines, respectively, in Figs 3b, 3c and 3d for 3 β values (20°, 40° and 60°).

With 2% tolerance for R_WS_, the calculations presented in Table E1 of supplement S2.E indicate that the approximations established in formulae (13) to (16) remain valid under the following condition:

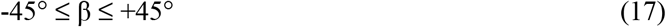

This specification will be required for the realization of a WS with the modalities determined by the data in Table 1.

### Linear relationship between the displacement of a hs and the rotation of the lever belonging to a head in working stroke

By arbitrarily matching the zero of the abscissa X with the zero of the angular positions θ and after integrating equation (15), the relationship between the abscissa (X_lin_; lin for *linear*) and θ is written:

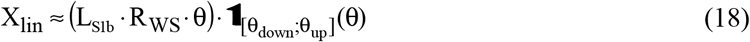

where 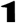 is the indicator function defined in (A2b) in Supplement S1.A of Paper 1.

With the values of θ_down_ and θ_up_ fixed in (12a) and (12b) and by respecting the conditions dictated in (17), the adequacy between the X_eq_ and X_lin_ abscissa formulated according to (11) and (18) is verified by linear regression for the 141 values of θ varying between θ_down_ and θ_up_ with a 0.5° pitch. The determination coefficient is higher than 99.9% in all cases; see Figs 4a, 4b and 4c where the X_eq_ and X_lin_ abscissa are represented as a function of θ by a pink and black line, respectively, for 3 values of the angle β.

**Fig 4.**
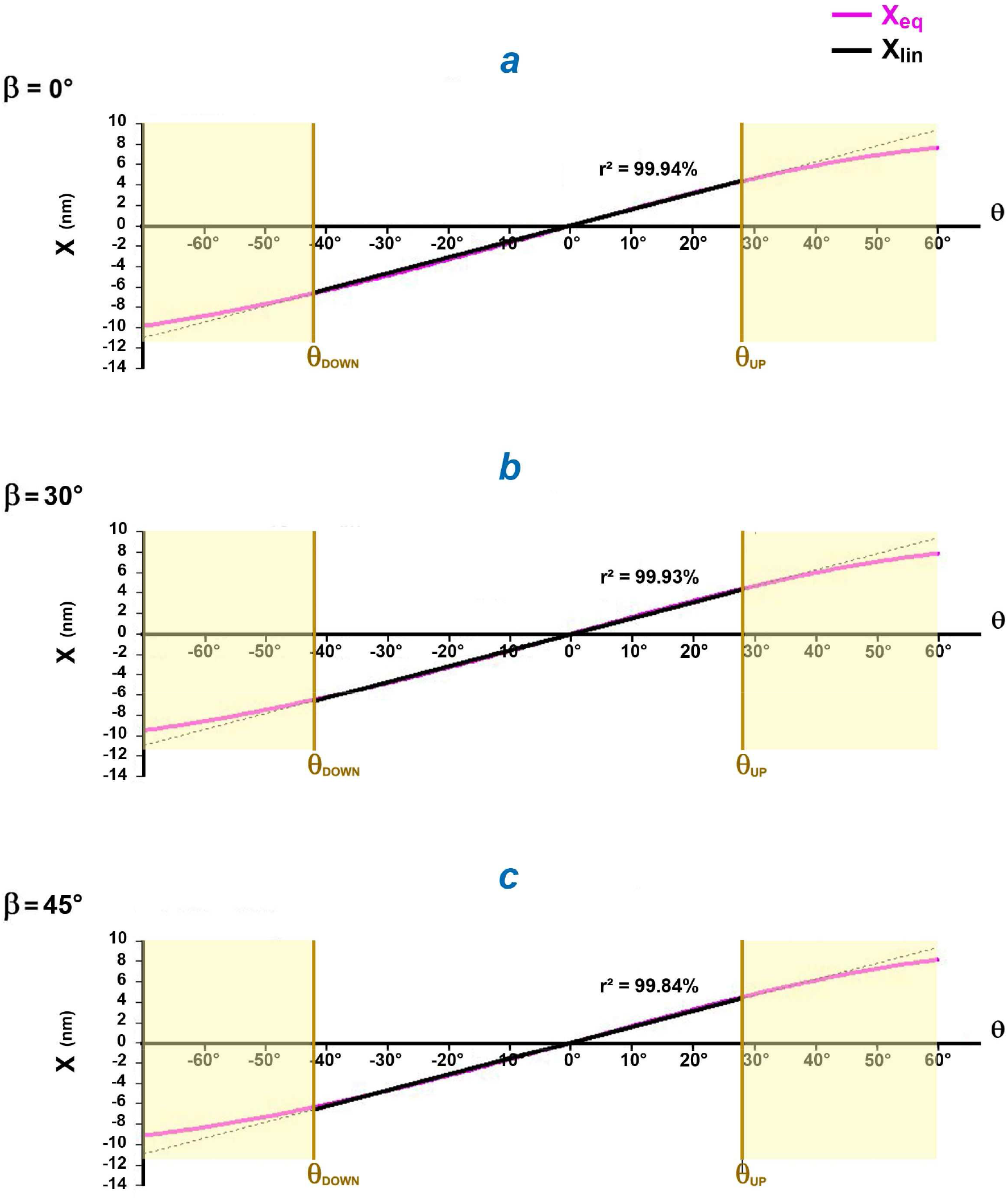
Evolutions of X_eq_ and X_lin_ abscissa in function of θ varying between −70 and +60° according to 3 values of β with α=0°. The determination coefficient (r^2^) is calculated by linear regression of X_eq_ as a function of X_lin_ between θ_down_ and θ_up_. (a) β = 0°. (b) β = 30°. (c) β = 45°.

From (18) we derive the linear relationship between the algebraic displacement of the hs (ΔX) and the corresponding rotation of the lever (Δθ) defined as:

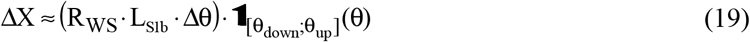

### Maximum step of one myosin head during WS (stroke size)

The maximum shortening of the hs (δX_Max_) corresponding to δθ_Max_ is called “stroke size”. It is calculated according to the linear approximation provided in (19):

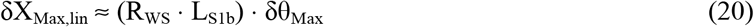

Note that the lever arm is not equal to L_S1b_ but to (R_WS_ · L_S1b_).

For work on intact vertebrate fibers, the stroke size generally varies between 10 and 12 nm [4,5,15,16,17,18,19]. The mean value of 11.5 nm is used for δX_Max_ confirmed by numerical application with the R_WS_ value determined in (13) and the data from Table 1:

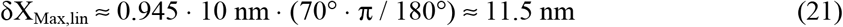

All tables in Supplement S2.E display in columns 4 and 5 the maximum steps δX_Max,eq_ and δX_Max,lin_ calculated according to relations (10) and (20), respectively. In all cases with the condition imposed in (17), the difference between the calculations of the 2 steps does not exceed 2%.

From now on, the stroke size (δX_Max_) will be calculated according to (20).

### Influence of the algorithmic method

With the zero slope method (see Methods section) used in paragraph E.2 of Supplement S2.E, there is a +1° difference compared to the values determined in (12a) and (12b) with the deviation minimization method. The data in Table E2 lead to conclusions similar to those in the previous paragraph. For all cases tested under identical conditions, the zero slope method provides similar results to the minimization of deviations, the difference for the values of de θ_down_ and θ_up_ never exceeding 1°.

### Negligible influence of α

The angle α varies between −30° and +30°; see expression (G5) of Supplement S3.G of accompanying Paper 3. The calculations made in Tables E3a and E3b of section E.3 in Supplement S2.E show that the value of α does not modify the conclusions indicated previously.

### Influence of lever length

The angles θ_down_ and θ_up_ are estimated with a standard L_S1b_ value equal to 10 nm [20.21]. In paragraph E.4 of Supplement S2.E, two other L_S1b_ values are tested: calculations with these 2 data yield deviations of ± 0.5° for θ_down_ and θ_up_. The reckonings exposed in Tables E4a and E4b induce conclusions consistent with those obtained when L_S1b_ is equal to 10 nm. The main difference is in the stroke size (δX_Max_) assessed at 11 and 12 nm for L_S1b_ equal to 9.5 and 10.4 nm, respectively. These results are logically ensued from the equality (20).

### Significant influence of inter-filament spacing (lattice)

In the previous calculations, the distance between 2 myosin filaments (d_2Mfil_) is equal to 46 nm (Table 1) as provided for the standard value in a skeletal fiber [7.8]. In paragraph E.5 of Supplement S2.E, two other d_2Mfil_ values were tested, 42 and 50 nm, for which respective deviations of +3° and −3° are noted with respect to the data seen in (11) and (12) for θ_down_ and θ_up_. There is a marked influence of the lattice on the θ angle in accordance with experimental observations: see Fig 8 in [22] and Fig 4 in [23] where the temperature rise increases the inter-filament spacing.

In conclusion, to take into account the variability of the data and methods of calculations, the values of θ_down_ and θ_up_ are specified with a common variability of ±5°.

## Discussion

### Hypothesis of the lever moving in a fixed plane during the Working Stroke

Extensive crystallographic and electron microscopy observations of the movement of the myosin head carried out *in situ* will reveal the validity of this assumption. However, there are concrete elements to support this hypothesis.

The idea that the movement of the myosin head is achieved in the same plane and therefore in a single direction is suggested on page 63 of the article princeps from I. Rayment [1] : « *However, the structure of the myosin head suggests that the power stroke arises from the reversal of domain movement in the myosin heavy chain induced by nucleotide binding* … *An immediate suggestion is that myosin forms a tight interaction with actin in only one orientation.* »

The article by A. Houdusse [11] evokes this point : « *The converter rotates about 65° between the transition and nucleotide-free states of scallop S1 and leads to a movement of the lever arm between these states that has a very small azimuthal component.* »

S. Hopkins’ work [24] reinforces the hypothesis, as an excerpt from the abstract indicates: « *We applied rapid length steps to perturb the orientations of the population of myosin heads that are attached to actin, and thereby characterized the motions of these force bearing myosin heads. During active contraction, this population is a small fraction of the total. When the filaments slide in the shortening direction in active contraction, the long axis of LCD tilts towards its nucleotide-free orientation with no significant twisting around this axis. In contrast, filament sliding in rigor produces coordinated tilting and twisting motions.* »

Or Arakelian’s [25] as suggested by an excerpt from the summary: « *We hypothesized that an azimuthal reorientation of the myosin motor domain on actin during the weak-binding to strong-binding transition could explain the lever arm slew provided that myosin’s α-helical coiled-coil subfragment 2 (S2) domain emerged from the thick filament backbone at a particular location*. »

If in our model the point A of fixing S1a on the surface of the actin molecule is unique, S1a is in reality strongly linked at several points [26,27,28,29], i.e. a precise and identical orientation for each S1a of a WS head towards the longitudinal axis of the Afil. Several studies [27,28] indicate a rotation of the mechanically guided lever S1b within the converter belonging to the motor domain, which again implies a single orientation.

In paragraph F.4 of Supplement S2.F, a mechanical study relating to the set of the WS myosin heads in a muscle fiber leads to two formulations of the force exerted at the ends of a myofibril, one by applying the linear and angular moment principles, the other according to the work-energy theorem. The equality between the 2 terms R_WS_ and S_WS_ delivered in (14) and the collinearity along the Oz axis of the vectors “moment” and “S1b angular rotation speed” for each moving WS head are essential so that the 2 expressions are equal. Precisely the hypothesis 4 of the lever moving in a fixed plane during the WS brings these 2 conditions.

### Corollaries of the hypothesis

The double helix structure of the Afil (see Supplement S3.G to Paper 3) prevents the two heads of a myosin molecule from being simultaneously in WS state. Indeed the two heads being attached to two distinct actin molecules, their respective levers would each evolve with a different angle β. However, S2 being the common rod linking at the Mfil, it is geometrically impossible that the two angles β remain constant when the Afil moves relative to the Mfil along the longitudinal axis OX. The corollary is documented by several researchers [30,31].

The equality (20) indicates a proportionality relationship between the stroke size (δX_Max_) and the length of the lever (L_S1b_,), a relationship observed experimentally [21,32]. Equality (20) is another relationship of proportionality between δX_Max_ and δθ_Max_, also interpreted [33].

## Conclusion

With the knowledge of the angles α, β and θ, the hypothesis of the lever displacement in a fixed plane allows the complete geometrization of the cross-bridge formed by a WS myosin head. Consequently, the relative position of the Mfil with respect to the Afil makes it possible to check if the WS state is feasible or not and in the confirmed case, to calculate the orientation θ of the lever. By applying this rule to an idealized half-sarcomere, it becomes possible to know the statistical distribution of θ. This is the subject of the study proposed in accompanying Paper 3.

## Supporting information

Supplementary Chapter. Kinematics and dynamics of a myosin head in Working Stroke.

Supplementary Chapter. Influence of various parameters on the 4 variables

Supplementary Chapter. Kinematics and dynamics of a stimulated muscle fiber and a myofibril.

Computer Programs for Paper 2

Data for Paper 2

Supplementary Chapter. Forces and energies involved in the working stroke of a myosin II head.

## Supporting information

### S2.C Supplementary Chapter. Forces and energies involved in the working stroke of a myosin II head

C.1 Units at the scale of a myosin head

C.2 Brownian displacements and thermal shocks

C.3 Hydrolysis of an ATP molecule and maximum tensile force of a myosin II head

C.4 Strong binding (SB)

C.5 Weak binding (WB)

C.6 Acceleration quantities of gravitational origin

C.7 Acceleration quantities of inertial origin

C.8 Archimedes’ Forces

C.9 Viscous type friction forces

C.10 Conclusion

References of Supplement S2.C

### S2.D Supplementary Chapter. Kinematics and dynamics of a myosin head in Working Stroke

D.1 Calculations relating to the kinematics of a WS myosin head with Fig D1

D.2 Calculations relating to the dynamics of a WS myosin head in a right hs shortening at constant velocity with Fig D2

### S2.E Supplementary Chapter. Influence of various parameters on the 4 variables R̅,S̅, δX_Max,eq_ and δX_Max,lin_

E.1 Influence of β displayed in Table E1

E.2 Influence of the algorithmic method displayed in Table E2

E.3 Influence of the angle α tested with α = +30° (Table E3a) and α = −30° (Table E3b)

E.4 Influence of the lever length with L_S1b_= 9.5 nm (Table E4a) and L_S1b_= 10.4 nm (Table E4b)

E.5 Influence of the inter-filament distance with d_2Mfil_ = 42 nm (Table E5a) and d_2Mfil_ = 50 nm (Table E5b)

### S2.F Supplementary Chapter. Kinematics and dynamics of a stimulated muscle fiber and a myofibril

F.1 Isolated fiber with Fig F1

F.2 Kinematics of a myofibril in the shortening or lengthening phase

F.3 Dynamics of a myofibril shortening or lengthening at constant speed with Fig F2

F.4 Theorem of work- energy applied to a muscle fiber shortening or lengthening at constant speed

References of Supplement S2.F

**CP2 Supplementary Material. Computer Programs for calculating the variables, X_eq_, X_lin_, R, S, res, and plotting the 5 variables as a function of θ**. Algorithms are written in Visual Basic 6.

**DA2 Supplementary Material. Data relating to computer programs used for Paper 2**. Access Tables are transferred to Excel sheets.

## References

1. Rayment I, Holden HM, Whittaker M, Yohn CB, Lorenz M, et al. (1993) Structure of the actin-myosin complex and its implications for muscle contraction. Science 261: 58–65.

2. Rayment I, Rypniewski WR, Schmidt-Base K, Smith R, Tomchick DR, et al. (1993) Three-dimensional structure of myosin subfragment-1: a molecular motor. Science 261: 50–58.

3. Baumketner A (2012) The mechanism of the converter domain rotation in the recovery stroke of myosin motor protein. Proteins 80: 2701–2710.

4. Ferenczi MA, Bershitsky SY, Koubassova N, Siththanandan V, Helsby WI, et al. (2005) The “roll and lock” mechanism of force generation in muscle. Structure 13: 131–141.

5. Geeves MA, Holmes KC (1999) Structural mechanism of muscle contraction. Annu Rev Biochem 68: 687–728.

6. Piazzesi G, Dolfi M, Brunello E, Fusi L, Reconditi M, et al. (2014) The myofilament elasticity and its effect on kinetics of force generation by the myosin motor. Arch Biochem Biophys 552-553: 108–116.

7. Millman BM (1998) The filament lattice of striated muscle. Physiol Rev 78: 359–391.

8. Squire JM, Al-Khayat HA, Knupp C, Luther PK (2005) Molecular architecture in muscle contractile assemblies. Adv Protein Chem 71: 17–87.

9. Geeves MA, Holmes KC (2005) The molecular mechanism of muscle contraction. Adv Protein Chem 71: 161–193.

10. Holmes KC, Geeves MA (2000) The structural basis of muscle contraction. Philos Trans R Soc Lond B Biol Sci 355: 419–431.

11. Houdusse A, Szent-Gyorgyi AG, Cohen C (2000) Three conformational states of scallop myosin S1. Proc Natl Acad Sci U S A 97: 11238–11243.

12. Irving M, Piazzesi G, Lucii L, Sun YB, Harford JJ, et al. (2000) Conformation of the myosin motor during force generation in skeletal muscle. Nat Struct Biol 7: 482–485.

13. Knowles AC, Ferguson RE, Brandmeier BD, Sun YB, Trentham DR, et al. (2008) Orientation of the essential light chain region of myosin in relaxed, active, and rigor muscle. Biophys J 95: 3882–3891.

14. Mello RN, Thomas DD (2012) Three distinct actin-attached structural states of myosin in muscle fibers. Biophys J 102: 1088–1096.

15. Piazzesi G, Reconditi M, Linari M, Lucii L, Sun YB, et al. (2002) Mechanism of force generation by myosin heads in skeletal muscle. Nature 415: 659–662.

16. Huxley H, Reconditi M, Stewart A, Irving T (2006) X-ray interference studies of crossbridge action in muscle contraction: evidence from muscles during steady shortening. J Mol Biol 363: 762–772.

17. Reconditi M, Linari M, Lucii L, Stewart A, Sun YB, et al. (2004) The myosin motor in muscle generates a smaller and slower working stroke at higher load. Nature 428: 578–581.

18. Piazzesi G, Lucii L, Lombardi V (2002) The size and the speed of the working stroke of muscle myosin and its dependence on the force. J Physiol 545: 145–151.

19. Goldman YE (1998) Wag the tail: structural dynamics of actomyosin. Cell 93: 1–4.

20. Reconditi M (2006) Recent Improvements in Small Angle X-Ray Diffraction for the Study of Muscle Physiology. Rep Prog Phys 69: 2709–2759.

21. Ruff C, Furch M, Brenner B, Manstein DJ, Meyhofer E (2001) Single-molecule tracking of myosins with genetically engineered amplifier domains. Nat Struct Biol 8: 226–229.

22. Colombini B, Bagni MA, Cecchi G, Griffiths PJ (2007) Effects of solution tonicity on crossbridge properties and myosin lever arm disposition in intact frog muscle fibres. J Physiol 578: 337–346.

23. Linari M, Brunello E, Reconditi M, Sun YB, Panine P, et al. (2005) The structural basis of the increase in isometric force production with temperature in frog skeletal muscle. J Physiol 567: 459–469.

24. Hopkins SC, Sabido-David C, van der Heide UA, Ferguson RE, Brandmeier BD, et al. (2002) Orientation changes of the myosin light chain domain during filament sliding in active and rigor muscle. J Mol Biol 318: 1275–1291.

25. Arakelian C, Warrington A, Winkler H, Perz-Edwards RJ, Reedy MK, et al. (2015) Myosin S2 origins track evolution of strong binding on actin by azimuthal rolling of motor domain. Biophys J 108: 1495–1502.

26. Holmes KC, Angert I, Kull FJ, Jahn W, Schroder RR (2003) Electron cryo-microscopy shows how strong binding of myosin to actin releases nucleotide. Nature 425: 423–427.

27. Kuhner S, Fischer S (2011) Structural mechanism of the ATP-induced dissociation of rigor myosin from actin. Proc Natl Acad Sci U S A 108: 7793–7798.

28. Behrmann E, Muller M, Penczek PA, Mannherz HG, Manstein DJ, et al. (2012) Structure of the rigor actin-tropomyosin-myosin complex. Cell 150: 327–338.

29. Mansson A, Rassier D, Tsiavaliaris G (2015) Poorly understood aspects of striated muscle contraction. Biomed Res Int 2015: 245154.

30. Juanhuix J, Bordas J, Campmany J, Svensson A, Bassford ML, et al. (2001) Axial disposition of myosin heads in isometrically contracting muscles. Biophys J 80: 1429–1441.

31. Brunello E, Reconditi M, Elangovan R, Linari M, Sun YB, et al. (2007) Skeletal muscle resists stretch by rapid binding of the second motor domain of myosin to actin. Proc Natl Acad Sci U S A 104: 20114–20119.

32. Uyeda TQ, Abramson PD, Spudich JA (1996) The neck region of the myosin motor domain acts as a lever arm to generate movement. Proc Natl Acad Sci U S A 93: 4459–4464.

33. Kohler D, Ruff C, Meyhofer E, Bahler M (2003) Different degrees of lever arm rotation control myosin step size. J Cell Biol 161: 237–241.

