## Supplementary Chapter. Kinematics and dynamics of a myosin head in Working Stroke. for "Mechanical model of muscle contraction 2. Kinematic and dynamic aspects of a myosin II head during the working stroke"

### S2.D Supplementary Chapter of Paper 2

### Kinematics and dynamics of a myosin II head in working stroke

#### D.1 Kinematics of a myosin head in working stoke (WS)

An actin filament (Afil) is modelled by a cylindrical rod with a radius $r_{Afil}$ oriented along $O_{Afil}X$, the main longitudinal axis where $O_{Afil}$ is an arbitrary point located on this axis (Fig D1b). A myosin filament (Mfil) is modelled by a cylindrical rod of radius $r_{Mfil}$ oriented according to $O_{Mfil}X$, the main longitudinal axis of the cylinder parallel to $O_{Afil}X$, where $O_{Mfil}$ is the orthogonal projection of $O_{Afil}$ on this axis. The $O_{Afil}Y°$ axis passing through the 2 points $O_{Afil}$ and $O_{Mfil}$ is perpendicular to $O_{Afil}X$ and the $O_{Afil}Z°$ axis is associated with $O_{Afil}Y°$ to define the transverse plane $O_{Afil}Y°Z°$ perpendicular to $O_{Afil}X$ (Fig D1b). The end of a myosin molecule is formed by 2 heads (Fig D1a). A myosin II head consists of two subfragments S1 and S2. The subfragment S1 is divided into two parts: the motor domain (S1a) and the lever (S1b) which are modelled by a cylinder and a cylindrical rod, respectively (Fig D1a). The two myosin heads are connected to the Mfil by a common rod, the S2 subfragment.

When S1a is strongly bound, the 5 segments, Afil, S1a, S1b, S2 and Mfil, form a poly-articulated chain commonly called as crossbridge. The 4 joints of the chain are represented by the 4 points A, B, C and D (Fig D1d). We examine the particular case where the segments S1a, S1b and S2 are rigid and where the points A, B and C are aligned with the $O_{Afil}$ point on the $O_{Afil}Y$ axis, which forms the angle β with $O_{Afil}Y°$ axis (Figs D1b and D1d). The $O_{Afil}Z$ axis is associated with $O_{Afil}Y$ to define the transverse plane $O_{Afil}YZ$ perpendicular to $O_{Afil}X$. Hypothesis n° 4 of the model states that the displacement of the BC segment, representing the lever S1b of a head in WS, is carried out in the fixed plane $O_{Afil}XY$, characterized by the constant angle β during any relative linear movement between the Afil and the Mfil (Figs D1b, D1c and D1d).

The 4 points, A°, B°, C° and D°, are the orthogonal projections of A, B, C and D on the $O_{Afil}XY°$ plane (Fig D1d).

Euclidean geometry provides the following two equations (Fig D1d):

$$d_{AMfil} = \left(r_{Afil} + AB_Y + L_{S1b} \cdot \cos\theta\right) \cdot \cos\beta - L^{o}_{S2} \cdot \sin\varphi° + r_{Mfil} \cdot \cos\alpha \qquad \text{(D1)}$$

$$L^{o}_{S2} = \sqrt{\left(L_{S2}\right)^2 - \left[\left(r_{Afil} + AB_Y + L_{S1b} \cdot \cos\theta\right) \cdot \sin\beta + r_{Mfil} \cdot \sin\alpha\right]^2} \qquad \text{(D2)}$$

where θ is the instantaneous angle between CB and CI in the $O_{Afil}XY$ plane such that point I is the projection of C on $O_{Afil}X$ according to $O_{Afil}Y$ (Fig D1c); α is the constant angle between $O_{Mfil}D°$ and $O_{Mfil}D$ ; β is the constant angle between $O_{Afil}Y°$ and $O_{Afill}Y$; φ° is the instantaneous angle between D°Y and D°C° in the $O_{Mfil}XY°$ plane; $AB_X$ and $AB_Y$ are the distances between A and B according to $O_{Afil}X$ and $O_{Afil}Y$ ; $d_{AMfil} = |O_{Afil}O_{Mfil}|$ ; $L_{S1b} = |BC|$ ; $L^{o}_{S2} = |C°D°|$ ; $L_{S2} = |CD|$.

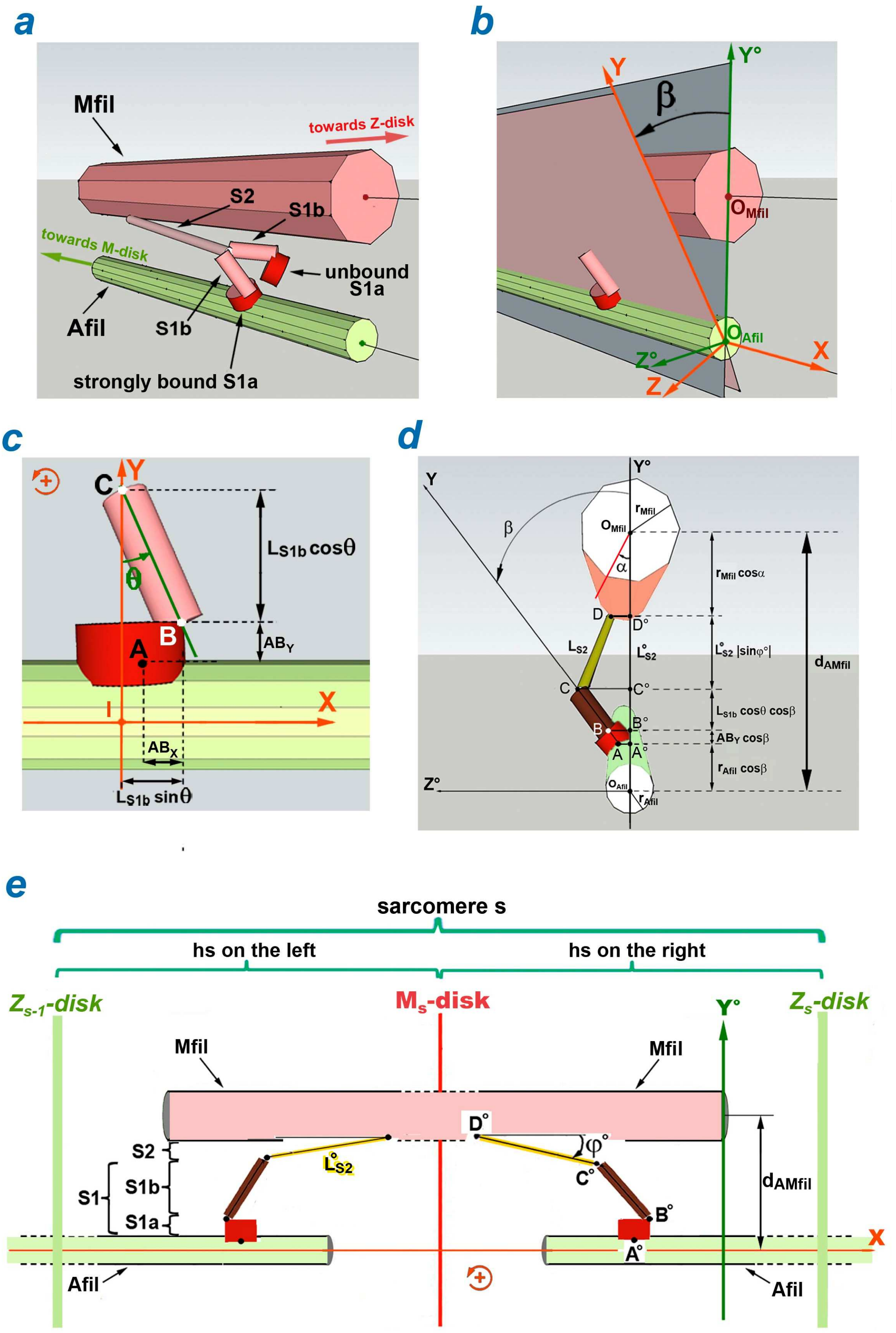


**Fig D1 Geometric characteristics of a WS myosin head.**

(a) Myosin molecule, one of whose 2 heads is in WS forming a cross-bridge between the myosin filament (Mfil) and the actin filament (Afil). (b) Definition of the angle β in the $O_{Afil}Y°Z°$ plane. (c) Definition of the θ angle in the IXY plane where point I is the projection of C on $O_{Afil}X$ according to $O_{Afil}Y$. (d) Cross-section in the $O_{Afil}Y°Z°$ plane where A°, B°, C° and D° are the orthogonal projections of A, B, C and D in the $O_{Afil}XY°$ plane. (e) Orthogonal projection of a schematized sarcomere in the $O_{Afil}XY°$ plane.

We fetch from the equality (D1) with (D2):

$$\varphi° = \mathrm{Arc}\sin\left(\frac{\left(r_{A\,fil} + AB_Y + L_{S1b}\cdot\cos\theta\right)\cdot\cos\beta + r_{Mfil}\cdot\cos\alpha - d_{AMfil}}{\sqrt{L_{S2}{}^{2} - \left[\left(r_{Afil} + AB_Y + L_{S1b}\cdot\cos\theta\right)\cdot\sin\beta + r_{Mfil}\cdot\sin\alpha\right]^{2}}}\right) \qquad \text{(D3)}$$

By definition, point A represents the recessed link between Afil and S1a, i.e. the binding site of the actin molecule, and D represents the ball joint between S2 and Mfil. In the $O_{Afil}XY°$ plane, we check (Figs D1c and D1e):

$$X_A = X_D - AB_X + L_{S1b}\cdot\sin\theta + L^{o}_{S2}\cdot\cos\varphi° \qquad \text{(D4)}$$

where $X_A$ and $X_D$ are the respective abscissa of A and D (or A° and D°) on the $O_{Afil}X$ and $O_{Mfil}X$ axes.

Arbitrarily, $X_A$ is imposed to cancel for $\theta$ =0, so:

$$X_D = AB_X - L^{o}_{S2}(\theta = 0)\cdot\cos\left[\varphi°(\theta = 0)\right]$$

With this condition, the abscissa $X_A$ is noted $X_{eq}$ and is reformulated:

$$X_{eq}(\theta) = L_{S1b}\cdot\sin\theta + L^{o}_{S2}(\theta)\cdot\cos\left[\varphi°(\theta)\right] - L^{o}_{S2}(\theta = 0)\cdot\cos\left[\varphi°(\theta = 0)\right] \qquad \text{(D5)}$$

The temporal derivation of equation (D2) provides:

$$\frac{dL^{o}_{S2}}{dt} = \dot{\theta}\cdot\frac{L_{S1b}\cdot\sin\beta\cdot\sin\theta\cdot\left[\left(r_{A\,fil} + AB_Y + L_{S1b}\cdot\cos\theta\right)\cdot\sin\beta + r_{Mfil}\cdot\sin\alpha\right]}{L^{o}_{S2}} \qquad \text{(D6)}$$

The derivation of (D1) gives with (D6):

$$0 = -\dot{\theta}\cdot\left(L_{S1b}\cdot\cos\beta\cdot\sin\theta\right) - \dot{\varphi}°\cdot\left(L^{o}_{S2}\cdot\cos\varphi°\right) - \dot{\theta}\cdot\sin\varphi°\cdot\frac{L_{S1b}\cdot\sin\beta\cdot\sin\theta\cdot\left[\left(r_{A\,fil} + AB_Y + L_{S1b}\cdot\cos\theta\right)\cdot\sin\beta + r_{Mfil}\cdot\sin\alpha\right]}{L^{o}_{S2}} \qquad \text{(D7)}$$

From (D7), it is extracted:

$$\dot{\varphi}° = -\dot{\theta}\cdot\frac{L_{S1b}\cdot\cos\beta\cdot\sin\theta}{L^{o}_{S2}\cdot\cos\varphi°} - \dot{\theta}\cdot\tan\varphi°\cdot\frac{L_{S1b}\cdot\sin\beta\cdot\sin\theta\cdot\left[\left(r_{A\,fil} + AB_Y + L_{S1b}\cdot\cos\theta\right)\cdot\sin\beta + r_{Mfil}\cdot\sin\alpha\right]}{\left(L^{o}_{S2}\right)^{2}} \qquad \text{(D8)}$$

The derivation of (D4) brings with (D6):

$$\dot{X}A = \dot{X}D + \dot{\theta}\cdot(L_{S1b}\cdot\cos\theta) - \dot{\varphi}^{\circ}\cdot\left(L^{o}_{S2}\cdot\sin\varphi^{\circ}\right) + \dot{\theta}\cdot\cos\varphi^{\circ}\cdot\frac{L_{S1b}\cdot\sin\beta\cdot\sin\theta\cdot\left[\left(r_{A\,fil}+AB_{Y}+L_{S1b}\cdot\cos\theta\right)\cdot\sin\beta + r_{Mfil}\cdot\sin\alpha\right]}{L^{o}_{S2}}$$

(D9)

It is posed:

$$res = \frac{\sin\beta\cdot\sin\theta\cdot\left[\left(r_{A\,fil}+AB_{Y}+L_{S1b}\cdot\cos\theta\right)\cdot\sin\beta + r_{Mfil}\cdot\sin\alpha\right]}{L^{o}_{S2}\cdot\cos\varphi^{\circ}} \qquad \text{(D10)}$$

Points A and D belong to the actin filament (Afil) and the myosin filament (Mfil), respectively. According to (D9) with (D8) and (D10), the relative shortening or lengthening velocity (u) of a hs on the right is equal to:

$$u = \dot{X}A - \dot{X}D = \dot{\theta}\cdot L_{S1b}\cdot\left[\left(\cos\theta + \sin\theta\cdot\tan\varphi^{\circ}\cdot\cos\beta\right) + res\right] \qquad \text{(D11)}$$

By symmetry of the sarcomere with respect to the M-disk, the expression (D11) is also valid for a hs on the left.

#### D.2 Dynamics of a WS head located in a hs on the right that shortens at constant speed

When a hs is shortened at a constant relative speed (u), all Afil and Mfil belonging to this hs move at a constant speed; see paragraph F.1 of Supplement S2.F. At the nanoscale, the quantities of linear and angular accelerations, of gravitational or inertial origin applied to the three segments S1a, S1b and S2, are nil or negligible; the same applies to the Archimedes' thrust and the viscosity forces acting on a myosin head; see Supplement S2.C. The only actions involved are the linking forces and moments. The pivotal motor-moment at point B between the 2 segments S1a and S1b is noted $\overrightarrow{\mathcal{M}_{B}}$ during the WS (Fig D2c). The linear moment principle (LMP) and the angular moment principle (AMP) are applied to the centers of gravity of the 3 non-deformable solids, S1a, S1b and S2, in the Galilean OXY°Z° reference frame, O being a fixed point in the laboratory, or a point located on Afil or Mfil.

We note $\gamma$ and $\delta$, respectively, the linear and angular accelerations of a solid.

#### Motor domain (S1a)

For the rigid segment S1a with $G_{S1a}$ as centre of gravity and $m_{S1a}$ as mass, the linking actions are modelled by a torsor (Figs D2a and D2b), whose reduction elements are $\left[\overrightarrow{F_A}\ ,\ -\overrightarrow{\mathcal{M}_A}\right]$ in A and $\left[-\overrightarrow{F_B}\ ,\ \overrightarrow{\mathcal{M}_B}\right]$ in B.

*LMP*

$$m_{S1a}\cdot\overrightarrow{\gamma_{G_{S1a}}} = \vec{0} = -\overrightarrow{F_B}\begin{vmatrix}-T_B\\ N_B\\ Q_B\end{vmatrix} + \overrightarrow{F_A}\begin{vmatrix}T_A\\ -N_A\\ -Q_A\end{vmatrix}$$

#### Lever (S1b)

For the rigid segment S1b with $G_{S1b}$ as centre of gravity and $m_{S1b}$ as mass, the linking actions are modelled by a torsor (Figs D2a, D2b and D2c), whose reduction elements are $\left[\overrightarrow{F_B}\ ,\ -\overrightarrow{\mathcal{M}_B}\right]$ in B and $\left[-\overrightarrow{F_C}\ ,\ \vec{0}\right]$ in C.

*LMP*

$$m_{S1b}\cdot\overrightarrow{\gamma_{G_{S1b}}} \approx \vec{0} = -\overrightarrow{F_C}\begin{vmatrix}-T_C\\ N_C\\ Q_C\end{vmatrix} + \overrightarrow{F_B}\begin{vmatrix}T_B\\ -N_B\\ -Q_B\end{vmatrix}$$

*AMP*

$$\overrightarrow{\delta_{G_{S1b}}} = \vec{0} = -\overrightarrow{\mathcal{M}_B}\begin{vmatrix}0\\ \mathcal{M}_B\cdot\sin\beta\\ -\mathcal{M}_B\cdot\cos\beta\end{vmatrix} + \overrightarrow{G_{S1b}B}\begin{vmatrix}\frac{L_{S1b}}{2}\cdot\sin\theta\\ -\frac{L_{S1b}}{2}\cdot\cos\theta\cdot\cos\beta\\ -\frac{L_{S1b}}{2}\cdot\cos\theta\cdot\cdot\sin\beta\end{vmatrix}\wedge\ \overrightarrow{F_B}\begin{vmatrix}T_B\\ -N_B\\ -Q_B\end{vmatrix} + \overrightarrow{G_{S1b}C}\begin{vmatrix}-\frac{L_{S1b}}{2}\cdot\cos\theta\\ \frac{L_{S1b}}{2}\cdot\cos\theta\cdot\cos\beta\\ \frac{L_{S1b}}{2}\cdot\cos\theta\cdot\sin\beta\end{vmatrix}\wedge\ -\overrightarrow{F_C}\begin{vmatrix}-T_C\\ N_C\\ Q_C\end{vmatrix}$$

#### Rod (S2)

For the rigid segment S2 with has $G_{S2}$ as centre of gravity and $m_{S2}$ as mass, the linking actions are modelled by a torsor (Figs D2a, D2b), whose reduction elements are $\left[\overrightarrow{F_C}\ ,\ \vec{0}\right]$ in C and $\left[-\overrightarrow{F_D}\ ,\ \vec{0}\right]$ in D.

*LMP*

$$m_{S2}\cdot\overrightarrow{\gamma_{G_{S2}}} \approx \vec{0} = -\overrightarrow{F_D}\begin{vmatrix}-T_D\\ N_D\\ Q_D\end{vmatrix} + \overrightarrow{F_C}\begin{vmatrix}T_C\\ -N_C\\ -Q_C\end{vmatrix}$$

*AMP*

$$\overrightarrow{\delta_{G_{S2}}} = \vec{0} = \overrightarrow{G_{S2}D}\begin{vmatrix}-\frac{\overset{\circ}{L}_{S2}}{2}\cdot\cos\varphi^\circ\\ -\frac{\overset{\circ}{L}_{S2}}{2}\cdot\sin\varphi^\circ\\ -\frac{L^z_{S2}}{2}\end{vmatrix}\wedge -\overrightarrow{F_D}\begin{vmatrix}-T_D\\ N_D\\ Q_D\end{vmatrix} + \overrightarrow{G_{S2}C}\begin{vmatrix}\frac{\overset{\circ}{L}_{S2}}{2}\cdot\cos\varphi^\circ\\ \frac{\overset{\circ}{L}_{S2}}{2}\cdot\sin\varphi^\circ\\ \frac{L^z_{S2}}{2}\end{vmatrix}\wedge\ \vec{F}\begin{vmatrix}T_C\\ -N_C\\ -Q_C\end{vmatrix}$$

where $L^z_{S2}$ is the orthogonal projection of $L_{S2}$ on the OZ° axis.

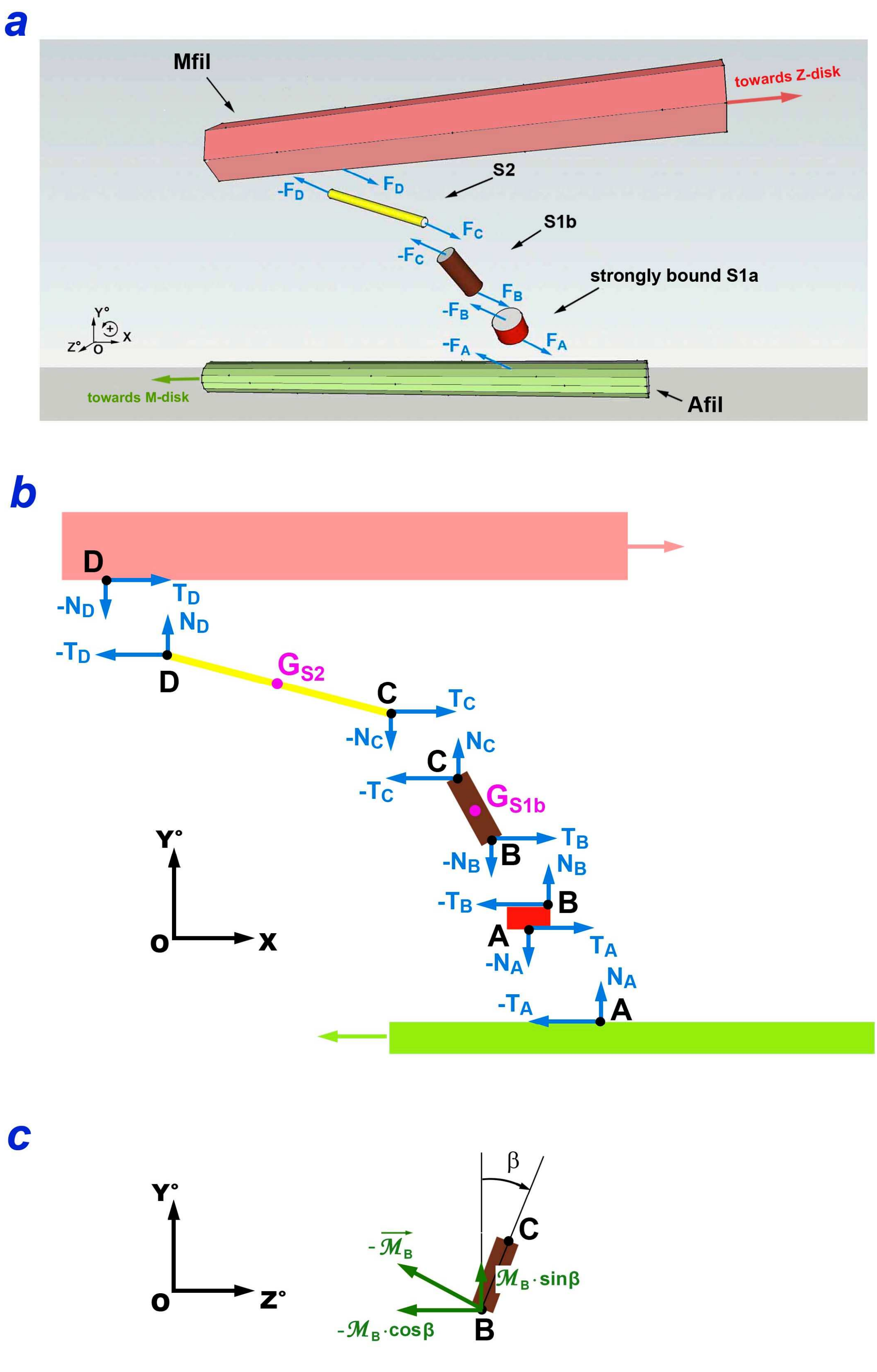


**Fig D2. Dynamic aspects of a WS myosin head.**

(a) Forces applied to the 5 rigid solids, Afil, S1a, S1b, S2, Mfil. (b) Orthogonal projections of these forces in the OXY° plane. (c) Moment $-\overrightarrow{\mathcal{M}_B}$ exerted in B on S1b according to OZ projected in the OY°Z° plane.

By noting that forces and moments are expressed in modulus, and angles in algebraic values, the 5 vectorial equations on the OX, OY° and OZ° axes provide the following algebraic equations:

**For S1a**

*On X:* $$0 = T_A - T_B \tag{D12}$$

*On Y°:* $$0 = -N_A + N_B \tag{D13}$$

**For S1b**

*On X:* $$0 = T_B - T_C \tag{D14}$$

*On Y°:* $$0 = -N_B + N_C \tag{D15}$$

*On Z°:* $$\mathcal{M}_B \cdot \cos\beta = \frac{L_{S1b}}{2} \cdot \left[ (T_B + T_C) \cdot \cos\beta \cdot \cos\theta - (N_B + N_C) \cdot \sin\theta \right] \tag{D16}$$

**For S2**

*On X:* $$0 = T_C - T_D \tag{D17}$$

*On Y°:* $$0 = -N_C + N_D \tag{D18}$$

*On Z°:* $$0 = (T_C + T_D) \cdot \sin\varphi° + (N_C + N_D) \cdot \cos\varphi° \tag{D19}$$

From (D12), (D14) and (D17), it is extracted:

$$T_A = T_B = T_C = T_D \tag{D20}$$

With (D13), (D15) and (D18), it is gotten:

$$N_A = N_B = N_C = N_D \tag{D21}$$

Equations (D16), (D19), (D20) and (D21) lead to:

$$\mathcal{M}_B = L_{S1b} \cdot T_A \cdot \left( \cos\theta + \frac{\sin\theta \cdot \tan\varphi°}{\cos\beta} \right) \tag{D22}$$
