## Supplementary Chapter. Influence of various parameters on the 4 variables for "Mechanical model of muscle contraction 2. Kinematic and dynamic aspects of a myosin II head during the working stroke"

### S2.E Supplementary chapter of Paper 2

### Influence of various parameters on the 4 variables, $\overline{\mathbf{R}}$, $\overline{\mathbf{S}}$, $\delta X_{Max,eq}$ and $\delta X_{Max,lin}$

**The geometric data below are constants for all calculations made in the 6 papers and related supplements:**

$r_{Afil}$ = 3.5 nm

$r_{Mfil}$ = 7.5 nm

$L_{S2}$ = 50 nm

$AB_X$ = 2 nm

$AB_Y$ = 1.5 nm

**The following geometric parameters are default values:**

$\alpha$ = 0°

$\beta$ = 0°

$d_{2Mfil}$ = 46 nm

$d_{AMfil} = d_{2Mfil} \cdot \sqrt{3}/3$ = 26.6 nm

$L_{S1b}$ = 10 nm *(actually 9.95 nm)*

These values are subject to change (see column "Variability range" in Table 1 of Paper 2).

For each table presented in this supplement, the deviation minimization method is used to calculate the values of $\theta_{up}$ and $\theta_{down}$ (see Methods section in Paper 2), except in paragraph E.2 where the zero slope method is adopted. Once these two bounds have been determined, all calculations are made by incrementing $\theta$ between $\theta_{up}$ and $\theta_{down}$ with a pitch of 0.5°.

**All supplement tables are presented in the same way:**

**1st column:** **value of the angle β**

**2nd column:** **$\overline{R}$ is the average value of the function R(θ,α,β) formulated with equation (2) of Paper 2**

**3rd column:** **$\overline{S}$ is the average value of the function S(θ,α,β) formulated with equation (6) of Paper 2**

**4th column:** **$\delta X_{Max,eq}$ is the stroke size calculated from equality (10) of Paper 2**

**5th column:** **$\delta X_{Max,lin}$ is the stroke size calculated from equality (20) of Paper 2**

**6th column:** **$r^2$ is the determination coefficient for the linear regression between the $X_{eq}$ and $X_{lin}$ abscissa calculated from equation (D5) in paragraph D.1 of Supplement S2.D and from equation (18) of Paper 2, respectively.**

**Note**

***Angles are presented in degrees but calculations are made in radian.***

***Any value that is outside its validity range is displayed in red.***

**Statistics**

The linear regression between $X_{eq}$ and $X_{lin}$ is graphically represented by a straight line passing through the origin, as the result:

$$X_{eq} = p \cdot X_{lin}$$

The slope (p) and the determination coefficient ($r^2$) are equal to:

$$p = \frac{\sum_{i=1}^{n} X_i \cdot Y_i}{\sum_{i=1}^{n} X_i^2}$$

$$r^2 = 1 - \frac{\sum_{i=1}^{n} (Y_i - p \cdot X_i)^2}{\sum_{i=1}^{n} \left(Y_i - \overline{Y}\right)^2}$$

### E. 1 Influence of the angle β

With α and β zero, the deviation minimization method provides:

$$\boldsymbol{\theta_{down} = -42°}$$

$$\boldsymbol{\theta_{up} = +28°}$$

Between these 2 terminals, the 4 parameters, $\overline{R}$, $\overline{S}$, $\delta X_{Max,eq}$, $\delta X_{Max,lin}$, are determined for 8 values of β displayed in the first column of Table E1. In the last column appears the determination coefficient relative to the linear regression between the 2 abscissa $X_{eq}$ et $X_{lin}$ between $\theta_{up}$ and $\theta_{down}$ with a pitch of 0.5°.

**Table E1. Calculations of 5 parameters for 8 values from β with the deviation minimization method and α=0°.**

| β | $\overline{R}$ | $\overline{S}$ | $\delta X_{Max,eq}$ (nm) | $\delta X_{Max,lin}$ (nm) | $r^2$ (%) |
|---|---|---|---|---|---|
| **0°** | **0.945** | **0.945** | 11.5 | 11.49 | 99.94 |
| **±10°** | 0.944 | 0.946 | 11.49 | 11.49 | 99.94 |
| **±20°** | 0.942 | 0.948 | 11.46 | 11.49 | 99.94 |
| **±30°** | 0.939 | 0.952 | 11.43 | 11.49 | 99.93 |
| **±40°** | 0.935 | 0.959 | 11.37 | 11.49 | 99.88 |
| **±45°** | 0.932 | 0.964 | 11.34 | 11.49 | 99.84 |
| **±50°** | 0.929 | 0.971 | 11.3 | 11.49 | 99.78 |
| **±60°** | 0.923 | 1.017 | 11.22 | 11.49 | 99.59 |

On the line β=0°, we find again the $R_{WS}$ and $S_{WS}$ values delivered to the equalities (13) of Paper 2.

If a tolerance threshold of 2% is set, a validity range of 0.926 to 0.969 is obtained for $R_{WS}$ and $S_{WS}$, and a validity range of 11.27 nm to 11.73 nm for $\delta X_{Max}$

From the reading of Table E1, it can be deduced that the linearity relationships provided in equations (15) and (16) of Paper 2 remain valid as long as β remains less than or equal to 45° in modulus.

We check:

$$\delta X_{Max,lin} = 0.945 \cdot 9.95 \text{ nm} \cdot (70° \cdot \pi / 180°) \approx 11.5 \text{ nm}.$$

### E.2 Influence of the algorithmic method

The zero slope method (see section Methods of Paper 2) is used to characterize the 2 bollards of θ with the result:

$$\theta_{down} = -41°$$

$$\theta_{up} = +29°$$

There is a difference of +1° compared to the values found in paragraph E.1. Between these two limits, the calculations of the variables studied for 8 values of β appear in Table E2.

**Table E2. Calculations of 5 parameters for 8 values from β with zero slope method and α=0°.**

| β | $\overline{R}$ | $\overline{S}$ | $\delta X_{Max,eq}$ (nm) | $\delta X_{Max,lin}$ (nm) | $r^2$ (%) |
|---|---|---|---|---|---|
| **0°** | 0.945 | 0.945 | 11.49 | 11.48 | 99.94 |
| **±10°** | 0.944 | 0.945 | 11.49 | 11.48 | 99.94 |
| **±20°** | 0.943 | 0.947 | 11.47 | 11.48 | 99.94 |
| **±30°** | 0.94 | 0.951 | 11.43 | 11.48 | 99.94 |
| **±40°** | 0.936 | 0.957 | 11.39 | 11.48 | 99.9 |
| **±45°** | 0.934 | 0.961 | 11.36 | 11.48 | 99.86 |
| **±50°** | 0.931 | 0.967 | 11.33 | 11.48 | 99.81 |
| **±60°** | 0.926 | 0.985 | 11.26 | 11.48 | 99.63 |

The data in Table E2 lead to the same conclusions compared to those in paragraph E.1.

### E.3 Influence of the angle α

#### Case a: α = +30°

With α = +30° and β = 0°, the deviation minimization method gives:

$$\theta_{down} = -42°$$
$$\theta_{up} = +28°$$

These are the same values determined with α=0° and the resulting calculations between these 2 bounds appear in Table E3a.

**Table E3a. Calculations of 5 parameters with α = +30°.**

| β | $\overline{R}$ | $\overline{S}$ | $\delta X_{Max,eq}$ (nm) | $\delta X_{Max,lin}$ (nm) | $r^2$ (%) |
|---|---|---|---|---|---|
| **0°** | 0.947 | 0.947 | 11.53 | 11.48 | 99.93 |
| **±10°** | 0.945 | 0.948 | 11.5 | 11.48 | 99.94 |
| **±20°** | 0.942 | 0.951 | 11.46 | 11.48 | 99.94 |
| **±30°** | 0.937 | 0.955 | 11.4 | 11.48 | 99.91 |
| **±40°** | 0.931 | 0.963 | 11.33 | 11.48 | 99.82 |
| **±45°** | 0.927 | 0.968 | 11.28 | 11.48 | 99.74 |
| **±50°** | 0.924 | 0.976 | 11.24 | 11.48 | 99.63 |
| **±60°** | 0.915 | 0.999 | 11.14 | 11.48 | 99.31 |

With a validity range of 0.926 to 0.969 for $R_{WS}$ and $S_{WS}$, and with a validity range of 11.27 nm to 11.73 nm for $\delta X_{Max}$, the values in Table E3a lead to the same conclusions compared to those in paragraph E.1.

**Case b: α= -30°**

With α = -30° and β = 0°, the deviation minimization method brings:

$$\theta_{down} = -42°$$

$$\theta_{up} = +28°$$

We find the values obtained with α=0° and the resulting calculations between these 2 terminals appear in Table E3b for 8 values from β.

**Table E3b. Calculations of 5 parameters with α = -30°.**

| β | $\overline{R}$ | $\overline{S}$ | $\delta X_{Max,eq}$ (nm) | $\delta X_{Max,lin}$ (nm) | $r^2$ (%) |
|---|---|---|---|---|---|
| **0°** | 0.947 | 0.947 | 11.53 | 11.48 | 99.93 |
| **±10°** | 0.948 | 0.948 | 11.53 | 11.48 | 99.92 |
| **±20°** | 0.947 | 0.95 | 11.53 | 11.48 | 99.93 |
| **±30°** | 0.945 | 0.955 | 11.5 | 11.48 | 99.94 |
| **±40°** | 0.942 | 0.962 | 11.46 | 11.48 | 99.94 |
| **±45°** | 0.94 | 0.967 | 11.44 | 11.48 | 99.93 |
| **±50°** | 0.938 | 0.974 | 11.41 | 11.48 | 99.91 |
| **±60°** | 0.932 | 0.995 | 11.33 | 11.48 | 99.83 |

With a validity range of 0.926 to 0.969 for $R_{WS}$ and $S_{WS}$, and with a validity range of 11.27 nm to 11.73 nm for $\delta X_{Max}$, the values in Table E3b lead to the same conclusions compared to those in paragraph E.1.

### E.4 Influence of lever length S1b ($L_{S1b}$)

**Case a: $L_{S1b}$ = 9.5 nm** *(actually 9.51 nm)*

With α and β zero, the deviation minimization method delivers:

$$\theta_{down} = -42.5°$$

$$\theta_{up} = +27.5°$$

There is a difference of -0.5° compared to the values found in paragraph E.1. Between these 2 bounds, the calculations of the variables studied for 8 values of β appear in Table E4a.

**Table E4a. Calculations of 5 parameters with $L_{S1b}$=9.5 nm.**

| **β** | $\bar{R}$ | $\bar{S}$ | $\delta X_{Max,eq}$ (nm) | $\delta X_{Max,lin}$ (nm) | $r^2$ (%) |
|---|---|---|---|---|---|
| **0°** | 0.946 | 0.946 | 11 | 11 | 99.94 |
| **±10°** | 0.946 | 0.947 | 11 | 11 | 99.94 |
| **±20°** | 0.945 | 0.949 | 10.97 | 11 | 99.94 |
| **±30°** | 0.943 | 0.953 | 10.93 | 11 | 99.93 |
| **±40°** | 0.94 | 0.961 | 10.87 | 11 | 99.88 |
| **±45°** | 0.932 | 0.966 | 10.84 | 11 | 99.85 |
| **±50°** | 0.929 | 0.973 | 10.8 | 11 | 99.79 |
| **±60°** | 0.922 | 0.996 | 10.72 | 11 | 99.61 |

It is noted that in this case

$$R_{WS} = S_{WS} = 0.946$$

$$\delta X_{Max,lin} = 0.946\ 9\cdot.5\ \text{nm}\ (\cdot 70°\ \cdot\pi/\ 180°) = 11\ \text{nm}.$$

With a tolerance threshold of 2%, we obtain a validity range between 0.927 and 0.965 for $R_{WS}$ and $S_{WS}$.

The $\bar{R}$ and $\bar{S}$ values in Table E4a lead to the same conclusions compared to those of paragraph E.1. However, for β=45°, the $S_{WS}$ value slightly exceeds the upper bound of the validity range.

**Case b: $L_{S1b}$ = 10.4 nm**

With α and β zero, the deviation minimization method gives:

$$\theta_{down} = -41.5°$$
$$\theta_{up} = +28.5°$$

There is a difference of +0.5° compared to the values found in paragraph E.1. Between these 2 limits, the calculations of the variables studied for 8 values of β appear in Table E4b.

**Table E4b. Calculations of 5 parameters with $L_{S1b}$= 10.4 nm.**

| B | $\overline{R}$ | $\overline{S}$ | $\delta X_{Max,eq}$ (nm) | $\delta X_{Max,lin}$ (nm) | $r^2$ (%) |
|---|---|---|---|---|---|
| **0°** | 0.944 | 0.944 | 12 | 12 | 99.94 |
| **±10°** | 0.943 | 0.945 | 12 | 12 | 99.94 |
| **±20°** | 0.942 | 0.947 | 11.98 | 12 | 99.94 |
| **±30°** | 0.939 | 0.951 | 11.94 | 12 | 99.93 |
| **±40°** | 0.936 | 0.957 | 11.89 | 12 | 99.88 |
| **±45°** | 0.932 | 0.962 | 11.85 | 12 | 99.84 |
| **±50°** | 0.929 | 0.968 | 11.82 | 12 | 99.77 |
| **±60°** | 0.923 | 0.988 | 11.74 | 12 | 99.58 |

It is noted that in this case

$$R_{WS} = S_{WS} = 0.944$$
$$\delta X_{Max,lin} = 0.944 \cdot 10.4\ \text{nm}\ (\cdot 70° \cdot \pi / 180°) = 12\ \text{nm}.$$

With a tolerance threshold of 2%, we obtain a validity range between 0.925 and 0.963 for $R_{WS}$ and $S_{WS}$.

The $\overline{\mathbf{R}}$ and $\overline{\mathbf{S}}$ values in Table E4b lead to the same conclusions compared to those of paragraph E.1.

### E.5 Influence of the inter-filament distance (*lattice*)

Reminder: the distance between 2 adjacent myosin filaments is noted as $d_{2Mfil}$ (see Table 1 of Paper 2)

#### Case a: $d_{2Mfil}$ = 42 nm (a decrease of 9% compared to 46 nm)

With α and β zero, the deviation minimization method brings**:**

$$\theta_{down} = -39°$$

$$\theta_{up} = +31°$$

There is a significant difference of +3° compared to the values found in paragraph E.1. Between these 2 boundaries, the calculations of the variables studied for 8 values of β appear in Table E5a.

**Table E5a. Calculations of 5 parameters with $d_{2Mfil}$=40 nm.**

| β | $\overline{R}$ | $\overline{S}$ | $\delta X_{Max,eq}$ (nm) | $\delta X_{Max,lin}$ (nm) | $r^2$ (%) |
|---|---|---|---|---|---|
| **0°** | 0.941 | 0.941 | 11.44 | 11.43 | 99.94 |
| **±10°** | 0.94 | 0.941 | 11.44 | 11.43 | 99.94 |
| **±20°** | 0.939 | 0.942 | 11.43 | 11.43 | 99.94 |
| **±30°** | 0.938 | 0.944 | 11.41 | 11.43 | 99.92 |
| **±40°** | 0.935 | 0.948 | 11.38 | 11.43 | 99.88 |
| **±45°** | 0.934 | 0.95 | 11.37 | 11.43 | 99.84 |
| **±50°** | 0.933 | 0.953 | 11.35 | 11.43 | 99.78 |
| **±60°** | 0.929 | 0.964 | 11.31 | 11.43 | 99.61 |

With a validity range between 0.926 and 0.969 for $R_{WS}$ and $S_{WS}$, it is verified that the $\overline{\mathbf{R}}$ and $\overline{\mathbf{S}}$ values in Table E5a all remain within the validity range and lead to different conclusions compared to those of paragraph E.1: the linearity relationships developed in equations (12) to (17) remain valid as long as β remains less than or equal to its maximum value of 60° in module, i.e. a value greater than 15° compared to the limit imposed for β in paragraph E.1.

#### Case b: $d_{2Mfil}$ = 50 nm (an increase of 9% compared to 46 nm)

With α and β zero, the deviation minimization method gives:

$$\theta_{down} = -45°$$

$$\theta_{up} = +25°$$

There is a difference of -3° compared to the values found in Paragraph E.1. Between these 2 extremes, the calculations of the variables studied for 8 values of β appear in Table E5a.

**Table E5b. Calculations of 5 parameters with $d_{2Mfil}$=50 nm.**

| β | $\overline{R}$ | $\overline{S}$ | $\delta X_{Max,eq}$ (nm) | $\delta X_{Max,lin}$ (nm) | $r^2$ (%) |
|---|---|---|---|---|---|
| **0°** | 0.951 | 0.951 | 11.58 | 11.57 | 99.93 |
| **±10°** | 0.95 | 0.953 | 11.56 | 11.57 | 99.93 |
| **±20°** | 0.947 | 0.956 | 11.53 | 11.57 | 99.92 |
| **±30°** | 0.942 | 0.963 | 11.46 | 11.57 | 99.92 |
| **±40°** | 0.935 | 0.974 | 11.38 | 11.57 | 99.88 |
| **±45°** | 0.931 | 0.983 | 11.33 | 11.57 | 99.84 |
| **±50°** | 0.926 | 0.994 | 11.27 | 11.57 | 99.78 |
| **±60°** | 0.916 | 1.028 | 11.14 | 11.57 | 99.58 |

With a validity range between 0.926 and 0.969 for $R_{WS}$ and $S_{WS}$ , it can be seen that the $\overline{\mathbf{R}}$ and $\overline{\mathbf{S}}$ values in Table E5b are outside the validity range as soon as β is equal to or greater than 35°, i.e. a value 10° lower than the limit imposed for β in paragraph E.1.
