## Supplementary Chapter. Kinematics and dynamics of a stimulated muscle fiber and a myofibril. for "Mechanical model of muscle contraction 2. Kinematic and dynamic aspects of a myosin II head during the working stroke"

### S2.F Supplementary Chapter of Paper 2

### Kinematics and dynamics of a stimulated muscle fiber and a myofibril

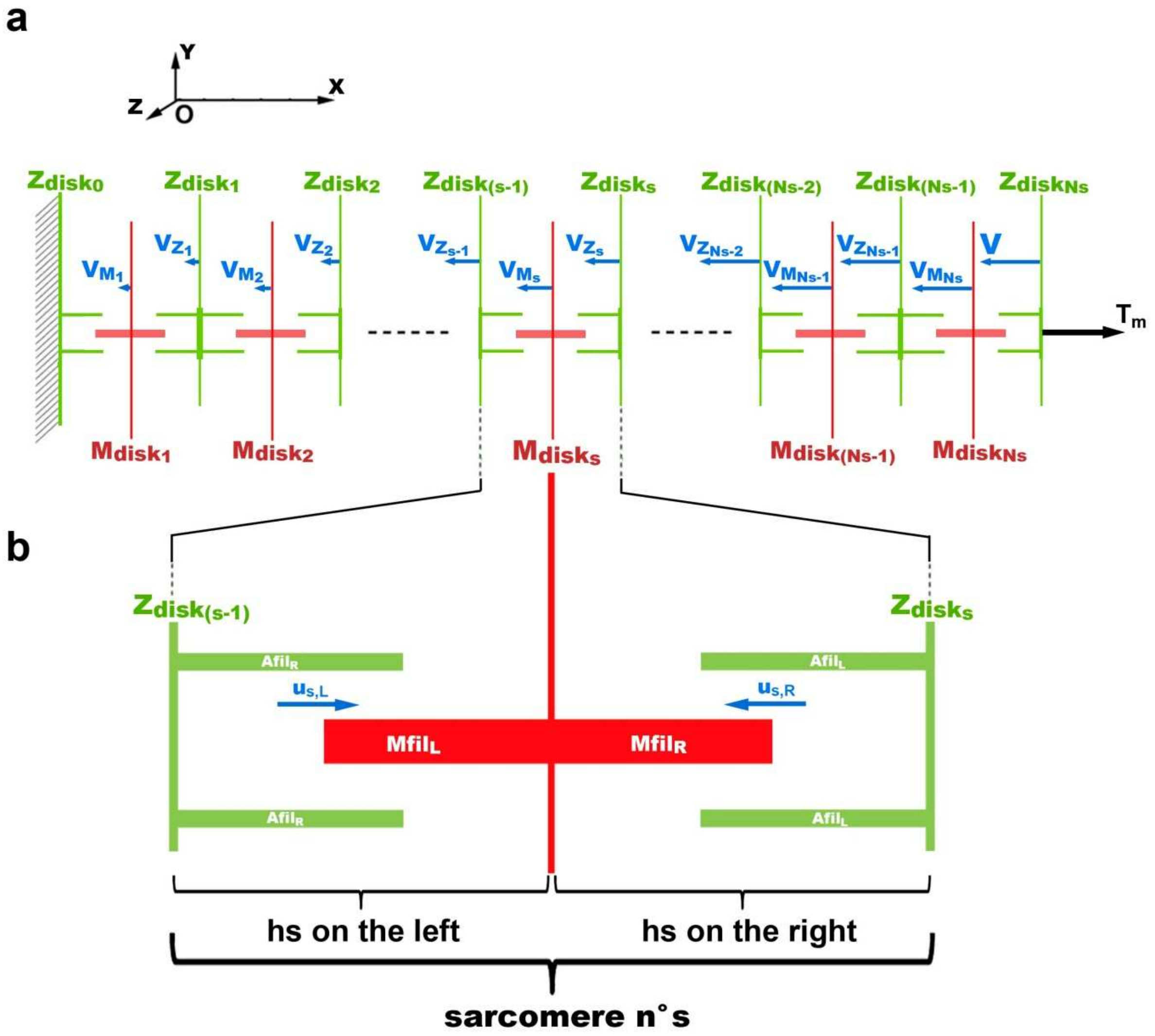


**Fig F1. Myofibril shortening at velocity V.**

(a) Myofibril composed of Ns sarcomeres arranged in series. (b) Sarcomere n° s composed of a half-sarcomere on the left and a half-sarcomere on the right shortening with the relative velocities, $u_{s,L}$ and $u_{s,R}$, respectively.

#### F.1 An isolated muscle fiber examined as a material system

A muscle fiber with its native structure is formed of $N_m$ identical myofibrils homogeneously distributed in the transverse plane Oyz, plane perpendicular to Ox, the permanent longitudinal axis of the fiber (Fig 1a). The fiber is prepared in such a way that its tendon structures play no role mechanically. During a shortening or lengthening phase at velocity V, the left end of the fiber remains fixed and a tension (T) is applied according to Ox to its free end on the right. Each myofibril changes its length at the same velocity V. Due to the homogeneity of the distribution, the force $T_m$ exerted at the end of each myofibril (Fig 1a) is equal to:

$$T_m = \frac{T}{N_m} \tag{F1}$$

Each myofibril consists of an identical number of sarcomeres (Ns) which are aligned in series according to Ox (Fig 1a). The sarcomere is the elementary component of the myofibril [1]. The sarcomere n° s, with s varying from 1 to Ns, is divided into two half-sarcomeres (Fig 1b):

1/ A half-sarcomere on the left (hsL) which consists of the transverse half of Z-disk n° (s-1) associated with all actin filaments ($AFil_R$) located to the right of Z-disk and the transverse half of M-disk n° s associated with all myosin filaments ($MFil_L$) located to the left of M-disk, such that:

$$hsL \equiv \left\{ 1/2 \cdot Zdisk_{s-1} + Afil_R \right\} \bigcup \left\{ 1/2 \cdot Mdisk_s + Mfil_L \right\} \tag{F2a}$$

where $\bigcup$ symbolizes the union of two sets.

2/ A half-sarcomere on the right (hsR) which consists of the transverse half of the M-disk n° s associated with all myosin filaments ($MFil_R$) located to the right of the M-disk and the transverse half of the Z-disk n° s associated with all myosin filaments located to the left ($AFil_L$) of the Z-disk, such that:

$$hsR \equiv \left\{ 1/2 \cdot Mdisk_s + Mfil_R \right\} \bigcup \left\{ 1/2 \cdot Zdisk_s + Afil_L \right\} \tag{F2b}$$

A myofibril is classically considered as the union of all right and left half-sarcomeres, i.e. the system called $\mathcal{S}1$ defined according to:

$$\mathcal{S}1 \equiv \bigcup_{i=1}^{Ns} \left\{ hsL_i \cup hsR_i \right\} \tag{F3}$$

If each sarcomere plays the same role whether it is located on the left or on the right, the myofibril is similar to a system, called $\mathcal{S}2$, made up of the union of all the undifferentiated half-sarcomeres:

$$\mathcal{S}2 \equiv \bigcup_{j=1}^{2\,Ns} \left\{ hs_j \right\} \tag{F4}$$

It is also possible to reconstruct a myofibril as the combination of all Z-disk associated with Afil located on the left and right (2Afil) and all M-disk associated with Mfil located on the right and left (2Mfil); this third system is named $\mathcal{S}3$ such that:

$$\mathcal{S}3 \equiv \{\text{Zdisk}_0 + \text{Afil}_R\} \cup \left( \bigcup_{k=1}^{Ns-1} \{\text{Zdisk}_k + 2\text{Afil}\} \right) \cup \{\text{Zdisk}_{Ns} + \text{Afil}_L\} \cup \left( \bigcup_{k=1}^{Ns} \{\text{Mdisk}_k + 2\text{Mfil}\} \right) \quad \text{(F5)}$$

The resting lengths of muscle fibers vary on average between 0.5 and 50 mm. The lengths of sarcomeres in vertebrate corresponding to these rest values are between 2 and 2.7 µm. It can be deduced from this that the number of sarcomeres (Ns) varies on average between 250 and 25 000.

#### F.2 Kinematics of a myofibril in the shortening or elongation phase

A myofibril is characterized according to the $\mathcal{S}3$ system proposed in (F5). During a shortening or lengthening of the fiber, all solids {Mdisk$_s$ + 2Mfil} or {Zdisk$_s$ + 2Afil}, are driven by linear translation movements according to Ox. Concerning the shortening of the sarcomere n° s (Fig 1b), we define $u_{s,L}$, the relative velocity of {Zdisk$_s$ + 2Afil} compared to {Mdisk$_s$ + 2Mfil}, and $u_{s,R}$, the relative velocity of {Zdisk$_s$ + 2Afil} compared to {Mdisk$_s$ + 2Mfil}.

We observe that $u_{s,L}$ is positive and $u_{s,R}$ negative in case of shortening of the myofibril. If the myofibril lengthens, the signs are reversed with $u_{s,L}$ negative and $u_{s,R}$ positive. Under isometric conditions, $u_{s,L}$ and $u_{s,R}$ are zero.

In the Galilean Oxyz reference frame of the laboratory, the velocity composition law gives by iteration the absolute velocity ( $V_{Z_s}$ ) of the solid {Zdisk$_s$ + 2Afil}:

$$V_{Z_s} = \sum_{j=1}^{s} \left( u_{j,R} - u_{j,L} \right) \quad \text{(F6)}$$

For s=Ns, we check:

$$V = V_{Z_{Ns}} = \sum_{j=1}^{Ns} \left( u_{j,R} - u_{j,L} \right) \quad \text{(F7)}$$

Similarly, we obtain the absolute velocity ( $V_{M_s}$ ) of the solid {Mdisk$_s$ + 2Mfil}, i.e. algebraically:

$$V_{M_s} = V_{Z_{s-1}} - u_{s,L} \quad \text{(F8)}$$

**<u>When a myofibril is shortened or lengthened at a constant speed V</u>**

If some non-deformable solids {$Mdisk_i$ + 2Mfil} or {$Zdisk_i$ + 2Afil} had accelerations, then these should be exactly compensated at each instant t by the decelerations of other rigid sets, {$Mdisk_j$ + 2Mfil} or {$Zdisk_j$ + 2Afil}, so that the constancy of V is ensured at all times. Kinetically, the acceleration or deceleration of a hs would be transmitted to neighbouring hs, generating fluctuations in V observed during critical oscillations [2] where the fiber is no longer in a stable state. Since the cycle of a cross-bridge is at the millisecond scale [2.3], the constancy of V observed for experiments lasting several hundred milliseconds requires the stability of the mechanical regime of the fiber. We deduce that absolute velocities $V_{M_s}$ and $V_{Z_s}$ are all constant.

From formulae (F6) to (F8), it is deduced by recurrence that the relative velocities $u_{s,L}$ and $u_{s,R}$ are all constant during the shortening or lengthening of the fiber achieved at constant V. We specify that this assertion does not mean that $us_{,L}$ and $_{us,R}$ are equal (see supplement S4.J of Paper 4).

Since sarcomeres behave identically for each of the $N_m$ myofibrils of the fiber, it is concluded from the above that the absolute velocities, $V_{M_s}$ and $V_{Z_s}$ and the relative velocities, $u_{s,L}$ and $u_{s,R}$, are identical with respect to the hsR or hsL n° s of any of the $N_m$ myofibrils.

#### Conclusion

With the equality (F1) and the previous remarks, the study of a myofibril is sufficient to understand the mechanical behaviour of the fiber when it is shortened or lengthened at a constant speed.

#### F.3 Dynamics of a myofibril shortening or lengthening at constant velocity

**Assessment of the forces acting on the constituent elements of a sarcomere**

The conclusion of Supplement S1.C states that at the nanoscale, the quantities of linear and angular accelerations, of inertial or gravitational origins, are negligible; the same applies to Archimedes' thrust and the viscosity forces acting on a myosin head. Thus the only mechanical actions present within a hs of the fiber are the linking forces and moments. Since the shortening movement occurs along the longitudinal axis Ox, only tangential forces are considered; radial forces, i.e. normal to the longitudinal axis of the myofibril, are statistically cancelled out by the geometric nature of the inter-filament space and the uniformity of the angular positions of the levers S1b belonging to the WS heads in each hs of the fiber (see Paper 3). For particular stretching or *rigor mortaris* conditions, the interaction forces of elastic origin caused by elongation, torsion or buckling of the S1b and S2 segments are grouped into a generic term called as Ti.

We introduce the viscosity forces exerted on the massive constituent elements of the sarcomere, i.e. {$Mdisk_s$ + 2Mfil} and {$Zdisk_s$ + 2Afil}. In this case, the moving velocity to be considered is the absolute speed of each of these components in relation to the laboratory's reference frame.

**Equation relative to a M-disk moving at constant speed**

A myofibril of the fiber is formed according to the $\mathcal{S}3$ system proposed in (F5). We consider the rigid set {$Mdisk_s$ + 2Mfil} distributed over the two hs located to the left and right of the sarcomere n° s (Figs F1b and F2). The actions applying to the solid {$Mdisk_s$ + 2Mfil} are the linking actions of the $\Lambda_{s,R}$ and $\Lambda_{s,L}$ myosin heads in WS, the interaction forces $-Ti_{M_{s,L}}$ and $Ti_{M_{s,R}}$ in case of stretching or *rigor mortaris*, and finally the viscosity forces $T_{Vi,M_s}$ .

The linear moment principle is implemented in the Oxyz reference frame of the laboratory to the non-deformable solid {$Mdisk_s$ + 2Mfil} moving at constant speed. The projection on OX of all the forces gives by summation (Fig F2):

$$0=\left(\sum_{b=1}^{\Lambda_{s,L}}\left(-T_{D_{s,L}^{(b)}}\right)+\sum_{b=1}^{\Lambda_{s,R}}\left(T_{D_{s,R}^{(b)}}\right)\right)+\left(-Ti_{M_{s,L}}+Ti_{M_{s,R}}\right)+T_{Vi,M_s} \qquad \text{(F9)}$$

where b is the index number of a WS head whose action causes at point $D_{s,L}^{(b)}$ ( $D_{s,R}^{(b)}$ ) an instantaneous force according to Ox equal to $-T_{D_{s,L}^{(b)}}$ $\left(T_{D_{s,R}^{(b)}}\right)$ whether the head is in hs on the left n° s (hs on the right n° s), respectively, and where point D is presented in paragraph D.2 of Supplement S2.D (Fig D2b).

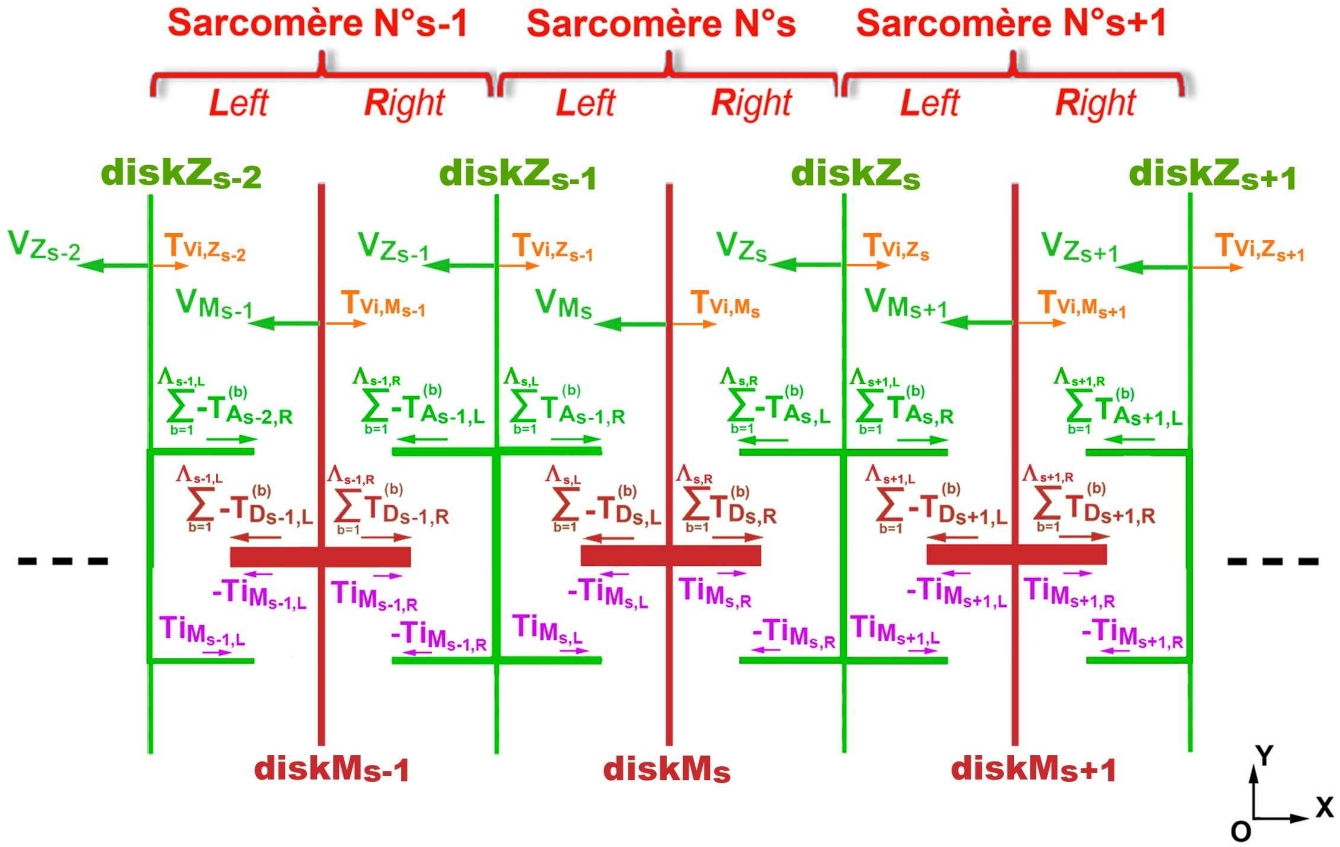


**Fig F2. Dynamics relative to a myofibril shortening at constant velocity.**

#### Equation relative to a Z-disk moving at constant speed

We consider the rigid solid {Zdisk$_s$ + 2Afil} distributed on the hs on right of the sarcomere n° s and the hs on left of the sarcomere n° (s+1) of a myofibril (Figs F1b and F2). The actions applying to {Zdisk$_s$ + 2Afil} are in the hs on the right of sarcomere n° s, the linking actions of $\Lambda_{s,R}$ heads in WS and the interaction forces $-Ti_{M_{s,R}}$ because equal in modulus and vectorially opposed to the actions applying to {Mdisk$_s$ + 2Mfil}, and in the hs on the left of the sarcomere n° (s+1) the linking actions of the $\Lambda_{s+1,L}$ heads in WS and the interaction forces $Ti_{M_{s+1,L}}$ because equal in modulus and vectorially opposed to the actions applying on {Mdisk$_{s+1}$ + 2Mfil}, and finally the viscosity forces $T_{Vi,Z_s}$.

The linear moment principle is implemented in the Oxyz reference frame of the laboratory to the non-deformable solid {Zdisk$_s$ + 2Afil} moving at constant speed. The projection on Ox of all the forces gives by summation (Fig F2):

$$0=\left(\sum_{b=1}^{\Lambda_{s,R}}\left(-T_{A^{(b)}_{s,L}}\right)+\sum_{b=1}^{\Lambda_{s+1,L}}\left(T_{A^{(b)}_{s,R}}\right)\right)+\left(-Ti_{M_{s,R}}+Ti_{M_{s+1,L}}\right)+T_{Vi,Z_s} \qquad \text{(F10)}$$

where b is the index number of a WS head whose action causes in $A^{(b)}_{s,L}$ ( $\mathbf{A^{(b)}_{s,R}}$ ) an instantaneous force according to Ox equal to whether $-T_{A^{(b)}_{s,L}}$ $\left(T_{A^{(b)}_{s,R}}\right)$ the head is in hs on the right n° s (hs on the left n° s+1), respectively; A is a point presented in paragraph D.2 of Supplement S2.D (Fig D2b).

For the last Z-disk n° Ns of the last sarcomere on which the tension $T_m$ is directly applied (Fig F1), the equation (F10) gives algebraically:

$$0=\sum_{b=1}^{\Lambda_{Ns,R}}\left(-T_{A^{(b)}_{Ns,L}}\right)+T_m-Ti_{M_{Ns,R}}+T_{Vi,Z_{Ns}} \qquad \text{(F11)}$$

Equations (F9), (F10) and (F11) are valid at any time t provided that the absolute velocity V is constant with negative for a shortening (or zero in isometry). In the case of a lengthening of the fiber at a constant and positive V velocity, the signs of the different forces are reversed in all three equations.

**Special case where the only actions involved are the linking forces and moments**

Under these conditions where the fiber shortens at a constant and slow velocity, the interaction forces and actions caused by viscosity are zero and equations (F9), (F10) and (F11) associated with the equality (4) of Paper 2 lead, iteratively and in modulus, to the following equations:

$$\sum_{b=1}^{\Lambda_{1,L}} T_{A_{0,R}^{(b)}} = \sum_{b=1}^{\Lambda_{1,L}} T_{D_{1,L}^{(b)}} = \sum_{b=1}^{\Lambda_{1,R}} T_{D_{1,R}^{(b)}} = \sum_{b=1}^{\Lambda_{1,R}} T_{A_{1,L}^{(b)}} = \sum_{b=1}^{\Lambda_{2,L}} T_{A_{1,R}^{(b)}} = \sum_{b=1}^{\Lambda_{2,L}} T_{D_{2,L}^{(b)}} = \sum_{b=1}^{\Lambda_{2,R}} T_{D_{2,R}^{(b)}} = \dots$$

$$\dots = \sum_{b=1}^{\Lambda_{s,L}} T_{A_{s-1,R}^{(b)}} = \sum_{b=1}^{\Lambda_{s,L}} T_{D_{s,L}^{(b)}} = \sum_{b=1}^{\Lambda_{s,R}} T_{D_{s,R}^{(b)}} = \sum_{b=1}^{\Lambda_{s,R}} T_{A_{s,L}^{(b)}} = \sum_{b=1}^{\Lambda_{s+1,L}} T_{A_{s,R}^{(b)}} = \sum_{b=1}^{\Lambda_{s+1,L}} T_{D_{s+1,L}^{(b)}} = \dots$$

$$\dots = \sum_{b=1}^{\Lambda_{Ns,L}} T_{A_{Ns-1,R}^{(b)}} = \sum_{b=1}^{\Lambda_{Ns,L}} T_{D_{Ns,L}^{(b)}} = \sum_{b=1}^{\Lambda_{Ns,R}} T_{D_{Ns,R}^{(b)}} = \sum_{b=1}^{\Lambda_{Ns,R}} T_{A_{Ns,L}^{(b)}} = T_m$$

(F12)

The tensions applied across any M or Z disks of a hs of the fiber are all equal in modulus when shortening or lengthening the myofibril at a constant speed. The series of equations displayed in (F12), which must be checked at all times, indicates that the number of WS heads is identical and that the spatial density of the angular positions of the levers belonging to these same heads is common in each hs of the fiber.

With a myofibril characterized by the $\mathcal{S}2$ system proposed in (F4), the equality series proposed in (F12) is rewritten in a more concise way for each hs of the fiber:

$$T_m = \sum_{b=1}^{\Lambda} T_D^{(b)} = \sum_{b=1}^{\Lambda} T_A^{(b)} \quad \text{(F13)}$$

where $\Lambda$ is the identical instantaneous number of myosin heads in WS per hs.

In support of the equalities (4) and (16) of Paper 2, (F13) becomes for each hs of the fiber:

$$T_m = \frac{1}{L_{S1b} \cdot S_{WS}} \cdot \sum_{b=1}^{\Lambda} \left| \mathcal{M}^{(b)} \right| \quad \text{(F14)}$$

where $L_{S1b}$ is the lever length; $S_{WS}$ is a constant calculated in equations (13) and (14) of Paper 2; $\mathcal{M}^{(b)}$ is the instantaneous motor moment generated by the head in WS n° b.

**Special case where the only actions involved are the linking forces and moments and where the tension T is constant**

We are in the presence of a stationary state of the fiber where each of the $N_m$ myofibrils shortens at constant speed V with constant $T_m$. The stability of this steady state is formulated according to the previous conclusions:

$$\Lambda(t+dt)=\Lambda(t)=\Lambda \qquad \text{(F15)}$$

$$f\left[\theta(t+dt)\right]=f\left[\theta(t)\right] \qquad \text{(F16)}$$

where Λ(t) is the number of myosin heads in WS per hs at time t; Λ is a constant number characteristic of the stationary state of the fiber; f is a function representing the spatial distribution of the θ angle between $\theta_{up}$ and $\theta_{down}$ identical in each hs.

This case is performed during the tetanus plateau under isometric conditions (V=0). In accompanying Paper 3, where a muscle fiber is studied under isometric conditions, the equations (F15) and (F16) are found again by geometric considerations, the density f present in (F16) being assumed as the Uniform law. This case also corresponds to phase 4 which follows a perturbation by a force step T where the muscle fiber shortens at constant speed.

It should be noted that equations (F12) to (F16) are implicitly formulated in A.F. Huxley's founding work [5].

#### F.4 The work-energy theorem applied to the fiber shortening at constant velocity

The work-energy theorem is written in power, *i.e.* in differential form:

$$\frac{dK_{fiber}}{dt}=P_{Fext}+P_{Fint} \qquad \text{(F17)}$$

where $K_{fiber}$ is the kinetic energy of the studied mechanical system, *i.e.* stimulated muscle fiber; $P_{Fext}$ is the power of external forces; $P_{Fint}$ is the power of internal forces.

The kinetic energy of a mechanical system is equal to the sum of the kinetic energies of translation and rotation of all the rigid bodies that compose it, i.e. in the case of fiber, all the solids {$Mdisk_s$ + 2Mfil} and {$Zdisk_s$ + 2Afil}, and in particular the rigid segments, S1a, S1b and S2 belonging to the myosin heads in WS. At a constant shortening velocity, as in phase 4 of any force step, these different non-deformable solids are all in uniform translation or rotation, with the exception of segments S1b and S2 where it is established that the inertial contributions of segments S1b and S2 are negligible; see paragraph C.7 of Supplement S1.C.

We deduce that the variation over time of the kinetic energy of a muscle fiber ($K_{fiber}$) shortening at constant speed is zero:

$$\frac{dK_{fiber}}{dt} = 0 \quad \text{(F18)}$$

By definition of the phase 4 of a force step, the fiber is subjected at one end to a constant tension T that works at constant speed V, and at the other fixed end to a tension T that does not work (Fig F1). The power of external forces is therefore formulated:

$$P_{Fext} = T \cdot V = -T \cdot |V| \quad \text{(F19)}$$

**<u>Special case where the only actions involved are the linking forces and moments and where the tension T is constant</u>**

In biomechanics or robotics, we can see that at the level of a joint that connects 2 rigid segments in motion, only the inter-segmental articular moment works. Also the power of a deformable mechanical system, which consists of rigid segments articulated between them and which moves in space, is equal to the sum of the powers of the articular moments. By applying this principle to the fiber being shortened at constant speed, the power of the internal forces of the fiber is equal to the sum of the powers of the articular moments of the WS heads. According to the hypothesis set out in Paper 2, the vectors of motor moments ($\mathcal{M}$) and angular velocities ( $\dot{\theta}$ ) are colinear along the Oz axis (Fig D2c in Supplement S2.D).

The power of the internal forces is formulated:

$$P_{Fint} = N_m \cdot \sum_{s=1}^{Ns} \left( \sum_{b=1}^{\Lambda_{s,L}} \dot{\theta}_{s,L}^{(b)} \cdot \mathcal{M}_{j,L}^{(b)} + \sum_{b=1}^{\Lambda_{s,R}} \dot{\theta}_{s,R}^{(b)} \cdot \mathcal{M}_{s,R}^{(b)} \right) \quad \text{(F20)}$$

where $N_m$ is the number of myofibrils; b is the index number of a WS head in the hs considered; $\Lambda_{s,L}$ and $\Lambda_{s,R}$ are, respectively, the number of WS heads in the hs located on the right and left of the sarcomere n° s of a myofibril.

By entering (F18), (F19) and (F20) in (F17), we obtain with (F1):

$$T_m = \frac{1}{|V|} \cdot \sum_{s=1}^{Ns} \left( \sum_{b=1}^{\Lambda_{s,L}} \dot{\theta}_{s,L}^{(b)} \cdot \mathcal{M}_{j,L}^{(b)} + \sum_{b=1}^{\Lambda_{s,R}} \dot{\theta}_{s,R}^{(b)} \cdot \mathcal{M}_{s,R}^{(b)} \right) \quad \text{(F21)}$$

With the equality (15) of Paper 2, (F21) is reformulated:

$$T_m = \frac{1}{L_{S1b} \cdot R_{WS} \cdot |V|} \cdot \sum_{s=1}^{Ns} \left( \sum_{b=1}^{\Lambda_{j,L}} u_{jsL} \cdot \mathcal{M}_{s,L}^{(b)} + \sum_{b=1}^{\Lambda_{s,R}} u_{s,R} \cdot \mathcal{M}_{s,R}^{(b)} \right) \tag{F22}$$

where $u_{s,L}$ and $u_{s,R}$ are, respectively, the relative linear velocities of the hs on the right and left of the sarcomere n° s, defined in Paragraph F.2.

Since the only actions involved are the linking forces and moments, equality (F14) applies and is rewritten with (F15) for each hs of the fiber:

$$T_m = \frac{1}{L_{S1b} \cdot S_{WS}} \cdot \sum_{b=1}^{\Lambda} \left| \mathcal{M}^{(b)} \right| \tag{F23}$$

where Λ is the constant number of heads in WS per hs.

The formulations given in (F22) and (F23) must remain unchanged and equal at any time **t**. This is the case here with (F15), (F16) and the two following conditions:

$$R_{WS} = S_{WS} \tag{F24}$$

$$\forall s, \; |u_{s,L}| = |u_{s,R}| = u = \frac{|V|}{2 \cdot N_S} \tag{F25}$$

Equality (F24) is verified in (14) in Paper 2 and equality (F25) is demonstrated: suppose that (F25) is not checked and that one hs shortens at a different speed than the other hs. To keep the temporal invariability of (F22), the number of WS heads and the distribution of θ of this hs must be modified and consequently the term for this hs no longer checks (F23).

In a stationary state of the muscle fiber where V and T are constant, the only solution that validates the equivalence between (F22) and (F23) is the one proposed with the equations (F15), (F16), (F24) and (F25).

#### Conclusion

In Paper 2, we assume that the lever of a WS head moves in a fixed plane containing the axis of the actin filament. The implication of this hypothesis on the inter-segmentary geometry of a WS head leads, on the one hand, to the expression (F22) established thanks to the colinearity of the vectors "moment" and "S1b angular rotation speed" for each WS head and, on the other hand, to equality (F24). The solution for the equivalence of (F22) and (F23) cannot be conceived without this hypothesis, thus providing an additional element to its validity.

#### References of Supplement S2.F to Paper 2


1. **Huxley AF (1957)** Muscle structure and theories of contraction. Prog Biophys Biophys Chem 7: 255-318.
2. **Huxley AF (1974)** Muscular contraction. J Physiol 243: 1-43.
3. **Ford LE, Huxley AF, Simmons RM (1974)** Proceedings: Mechanism of early tension recovery after a quick release in tetanized muscle fibres. J Physiol 240: 42P-43P.
4. **Lombardi V, Piazzesi G, Linari M (1992)** Rapid regeneration of the actin-myosin power stroke in contracting muscle. Nature 355: 638-641.
5. **Huxley HE (1957)** The double array of filaments in cross-striated muscle. J Biophys Biochem Cytol 3: 631-648.
