## Supplementary material for "Mechanical model of muscle contraction 2. Kinematic and dynamic aspects of a myosin II head during the working stroke": Computer Programs for Paper 2

### CP2 Computer programs for calculating the stroke size ($\delta X_{Max}$) and plotting R, S, res , $\delta X_{lin}$ and $\delta X_{eq}$ as a function of θ

All the characteristic data of a myosin head and a hs are displayed on the computer screen by starting the Sub PLANOUIEaffi() routine. The data are stored in an ACCESS table and can be manually modified (Fig CP2.1). As soon as a parameter is changed all calculations are performed again with sub-routine SUB_PLANOUIEcalculs().

By clicking on the orange button (Fig CP2.1), the different curves required for Paper 2 are drawn by starting the Sub VIT_raccsst_tetM() routine (Fig CP2.2 and CP2.3).

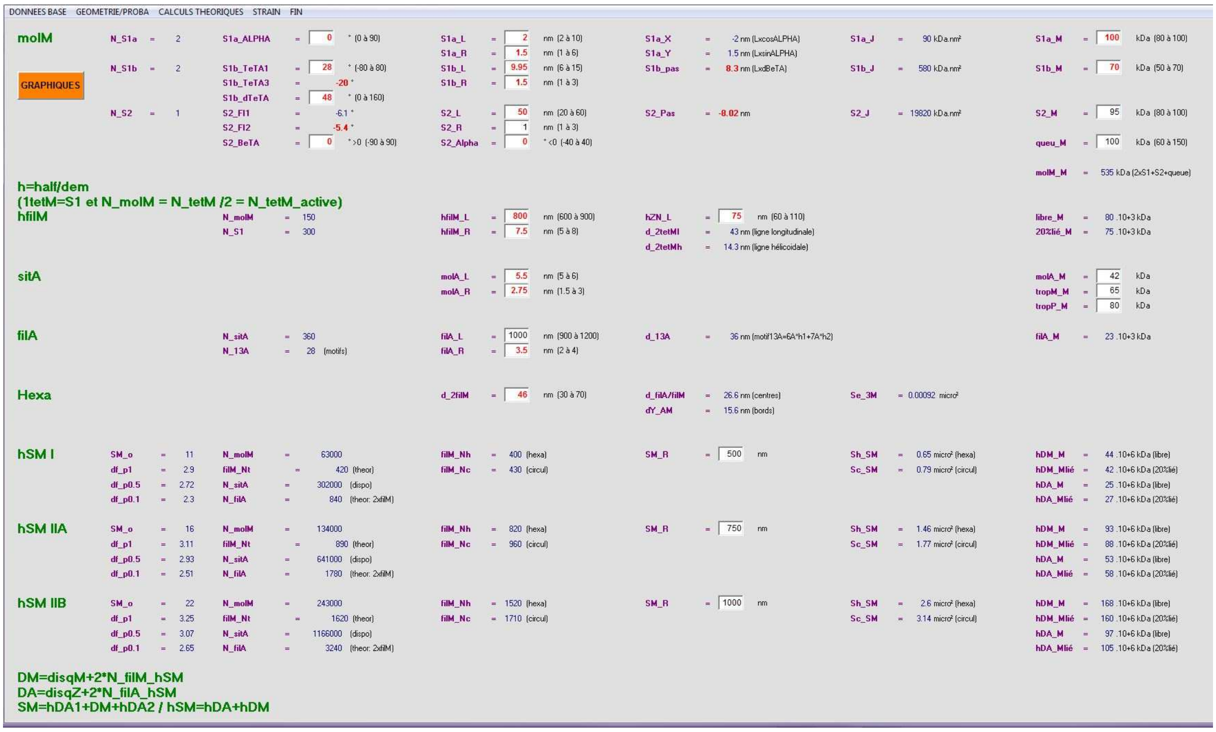


**Fig CP2.1. Screenshot after starting the "Sub AA_PLANOUIEaffi()" routine**

#### Sub PLANOUIEaffi()

```
'rappels: 1 molM = 2 tetM + S2(neck) = 2 S1 + S2 = 2(S1a+S1b)+S2
'   avec  1 tetM = S1 = S1a + S1b

Dim col As Integer, S1 As Integer, S2 As Integer, zki As String, zke As String, zkoi As String
sk.Index = "kikekoi"

If mmm = 1 Then
    Set X = Planouie
    X.Visible = -1
    X.Cls
    For t = 0 To N_Tx: Tx(t).Visible = 0: Next
    Set Cdrecalcul.Container = X
    With Cdrecalcul: .Caption = "RECAL CUL": .Tag = "Planouie": End With
    'Set Frlaser.Container = X 'X = Planouie
    'Set FrBIB.Container = X
    Frlaser.Visible = -1
    PUBMED.Visible = 0
Else 'mmm=2  impression laser ou PDF
    Set X = Printer
    'X.ForeColor = QBColor(0)
End If

With X

'li = -2 'compteur de lignes
yn = 2 'hauteur entre 2 lignes
'yt = 0 'compteur de saut de lignes pour le koi considéré
'yx = 0 'saut total pour le koi considéré


'principe: les données sont table_skonpeu et mise en page ds table_pages

pp.Seek ">=", zp, -10 'page +ligne 'en general commence àla ligne 0
If pp.NoMatch Then Stop

Do While pp("page") = zp
   For col = 0 To 6
    If IsNull(pp("col" + Format(col))) = 0 Then
        tx2 = Trim(pp("col" + Format(col)))

        'postionnement colonne
        'xx = debut enX pour colonne considérée
        xx = 0
        If col = 0 Then 'entete
            xx = 1
        Else
           'on cherche la variable' sk("col"), sk("entete") + "> " + sk("koi") + "> " + sk("lb")
           S1 = InStr(tx2, ">"): If S1 = 0 Then Stop
           zki = Mid(tx2, 1, S1 - 1)
           S2 = InStr(S1 + 1, tx2, ">"): If S2 = 0 Then Stop
           zke = Mid(tx2, S1 + 2, S2 - S1 - 2)
           zkoi = Mid(tx2, S2 + 2, Len(tx2) - S2)
           sk.Seek "=", zki, zke, zkoi: If sk.NoMatch Then Stop

           'xt = espacement avant signe egal et label
           'xn = espacement apres signe egal et valeur calculée (vv2)
```

```
  xn = 5.3: xt = 0 ': xn = 0

   'quelle colonne - on s'emmerde pas, on recrée une tabulation pour chaque page du menu
   Select Case zp
   Case 1
      Select Case col
       Case 1: xx = 8: xt = 0.8: xn = 5.5
       Case 2: xx = 16.5: xn = 7: xt = 2.5 'xn = 8.4: xt = 1
       Case 3: xx = 33: xt = 0.8
       Case 4: xx = 48.5: xt = 1.5: xn = 5.5
       Case 5: xx = 64: xt = 0.6: xn = 5.5
       Case 6: xx = 78: xt = 0.6
       Case Else: Stop ' pas normal'debug.print sk("i1");"-";sk("i2");"-";sk("i3")
      End Select
    Case 2
      Select Case col
       Case 1: xx = 8.3: xt = 2: xn = 5.5
       Case 2: xx = 20: xn = 7: xt = 3.2
       Case 3: xx = 40: xt = 3: xn = 6
       Case 4: xx = 57: xt = 1.5: xn = 5.5
       Case 5: xx = 64: xt = 1.5
       Case 6: xx = 78: xt = 1.5
       Case Else: Stop ' pas normal'debug.print sk("i1");"-";sk("i2");"-";sk("i3")
      End Select
    Case 3
     Select Case col
      Case 1: xx = 10: xt = 2: xn = 5.5
      Case 2: xx = 27: xn = 7: xt = 3.2
      Case 3: xx = 38: xt = 1.5
      Case 4: xx = 51: xt = 1.5: xn = 5.5
      Case 5: xx = 64: xt = 1.5
      Case 6: xx = 78: xt = 1.5
      Case Else: Stop ' pas normal'debug.print sk("i1");"-";sk("i2");"-";sk("i3")
     End Select
   End Select

End If
If xx = 0 Then Stop 'pas normal
 'quelle ligne
li = (pp("lig") + 1) * yn
.FontBold = -1

'*******************************
If xx = 1 Then 'en tete

   .FontSize = 14
   If mmm = 1 Then .ForeColor = QBColor(2)

   If tx2 = "Transv: 1 hSM" Then
      .CurrentX = xx + 13
      .CurrentY = 2
   ElseIf tx2 = "Longit: n SM/cm musc" Then
       .CurrentX = xx + 63
       .CurrentY = 2
   Else
      .CurrentX = xx
      .CurrentY = (pp("lig") + 0.7) * yn
   End If
```

```
    X.Print tx2
    '.FontBold = 0
    .FontSize = 9

'**********************************
ElseIf sk("ty") = 1 Then 'val. base
   If mmm = 1 Then .ForeColor = QBColor(5)
   .CurrentX = xx: .CurrentY = li
   If sk("lb") = "N" Or sk("lb") = "d" Then
       tx1 = sk("lb") + "_" + sk("koi")
   Else 'le reste
       tx1 = sk("koi") + "_" + sk("lb")
   End If
   X.Print tx1
   .CurrentX = xx + xt + 3: .CurrentY = li: X.Print "="
   If mmm = 1 Then 'on affiche la case
        With Tx(sk("no"))
         .Left = xx + xt + 4: .Top = li - 0.5: .Visible = -1
         If mmm = 1 Then .ForeColor = QBColor(IIf(sk("mm") = 1, 12, 0))
         .FontBold = IIf(sk("mm") = 1, -1, 0)
         .Text = Format(sk("vv1"))
        End With

   Else
      .CurrentX = xx + xt + 4: .CurrentY = li
      .FontBold = -1: .FontItalic = -1
      X.Print Format(sk("vv1"))
      .FontBold = 0: .FontItalic = 0
   End If

   'unité associée à la valeur affichée
   If IsNull(sk("uu")) = 0 Or IsNull(sk("uD")) = 0 Then
      .CurrentX = xx + xt + 7: .CurrentY = li - 0.1
      .ForeColor = QBColor(0): .FontBold = 0
      tx1 = ""
      If IsNull(sk("uD")) = 0 Then tx1 = IIf(IsNull(sk("uD")), "", "." + sk("uD") + " ")
      If IsNull(sk("uu")) = 0 Then tx1 = tx1 + sk("uu")
      If IsNull(sk("min")) = 0 Then tx1 = tx1 + "  (" + Format(sk("min")) + " à " + Format(sk("max")) + ")"
      X.Print tx1
   End If

'**********************
ElseIf sk("ty") = 2 Then 'val .calculée
   '.FontItalic = -1
   If mmm = 1 Then .ForeColor = QBColor(5)
   If sk("lb") = "N" Or Mid(sk("lb"), 1, 1) = "d" Or Mid(sk("lb"), 1, 1) = "S" Then
     tx1 = sk("lb") + "_" + sk("koi")
   Else 'le reste
      tx1 = sk("koi") + "_" + sk("lb")
   End If
   .CurrentX = xx: .CurrentY = li: X.Print tx1

   xav = IIf(sk("lb") = "N", 2.2, 3)
   .CurrentX = xx + xt + xav: .CurrentY = li: X.Print "="
   tx1 = Format(sk("vv2"))
   If mmm = 1 Then .ForeColor = QBColor(IIf(sk("mm") = 1, 12, 1))
   .FontBold = IIf(sk("mm") = 1, -1, 0)
```

```
          xap = IIf(sk("i1") < 10 Or sk("i1") = 31, 5, 7)
          If sk("lb") = "N" Then
           .CurrentX = xx + xt + xap - 0.5 - .TextWidth(tx1)
          ElseIf sk("lb") = "Se" Then
            .CurrentX = xx + xt + xap - 1.2
          Else
           .CurrentX = xx + xt + xn - .TextWidth(tx1)
          End If

          .CurrentY = li: X.Print tx1
          .ForeColor = QBColor(0)
          .FontBold = 0

        'unité associée à la valeur affichée + commentaire 'lb2' si il y en un
        If IsNull(sk("uu")) = 0 Or IsNull(sk("uD")) = 0 Or IsNull(sk("lb2")) = 0 Then
          If sk("lb") = "N" Then
             .CurrentX = xx + xt + xap
           ElseIf sk("lb") = "Se" Then
              .CurrentX = xx + xt + xap - 0.9 + .TextWidth(tx1)
          Else
              .CurrentX = xx + xt + xn + 0.2
          End If
          .CurrentY = li
          .ForeColor = QBColor(0)
          tx1 = ""
          If IsNull(sk("uD")) = 0 Then tx1 = IIf(IsNull(sk("uD")), "", "." + sk("uD") + " ")
          If IsNull(sk("uu")) = 0 Then tx1 = tx1 + sk("uu")
          If IsNull(sk("lb2")) = 0 Then tx1 = tx1 + " (" + sk("lb2") + ")"
          X.Print tx1

        End If
        '.FontItalic = 0
    End If

  End If 'len(tx2)>0
  Next 'col

  pp.MoveNext
  If pp.EOF() Then Exit Do
   'If sk("i1") > i1F Then
   '   Exit Do
   'ElseIf sk("i1") = 2 Or sk("i1") = 17 Then
   '  li = li + 1
   'ElseIf sk("i1") = 3 Then

  'ElseIf sk("i1") = 18 Or sk("i1") = 19 Or sk("i1") = 51 Then 'les regles pour hSM et SM
  '    li = 3
  'ElseIf (sk("i1") = 1 And sk("koi") <> vd) Or (sk("i1") > 1 And sk("entete") <> vd) Then
  '    li = li + yx + yn + 0.5
  '    yt = 0: yx = 0
  ' End If

Loop

End With
End Sub
```

#### Sub PLANOUIEcalculs()

```
 '** Position depart WORKING STROKE
   sk.Seek "=", "molM", "S2", "BeTA": BeTA = DEGenRD(sk("vv1"))
   sk.Seek "=", "molM", "S2", "Alpha": ALPHA_S2rd = DEGenRD(sk("vv1"))
   'LS2_p  : p=projection de LS2 ds plan OXY°
   LS2_p = Sqr(LS2 ^ 2 - ((r_filA + YS1a + LS1b * Cos(Teta1)) * Sin(BeTA) + r_filM * Sin(ALPHA_S2rd)) ^ 2)
   v = (r_filA + YS1a + LS1b * Cos(Teta1)) * Cos(BeTA) + r_filM * Cos(ALPHA_S2rd) - d_filAMc
   FI1 = ASINr(v / LS2_p) 'en rd
   X1 = LS1b * Sin(Teta1) + LS2_p * Cos(FI1)


  '** Position finale WORKING STROKE
   LS2_p = Sqr(LS2 ^ 2 - ((r_filA + YS1a + LS1b * Cos(Teta3)) * Sin(BeTA) + r_filM * Sin(ALPHA_S2rd)) ^ 2)
   v = (r_filA + YS1a + LS1b * Cos(Teta3)) * Cos(BeTA) + r_filM * Cos(ALPHA_S2rd) - d_filAMc
   FI2 = ASINr(v / LS2_p) 'en rd
   PLANOUIEvv2 "molM", "S2", "FI2", FI2 * 180 / pi, -2 'FI2 en °
   X3 = LS1b * Sin(Teta3) + LS2_p * Cos(FI2)
   PLANOUIEvv2 "molM", "S2", "Pas", X3 - X1, -2 'pas  tetM

End Sub
```

##### Function ASINr(xt As Single) As Single 'calcul avec angle en radian

```
 ASINr = xt + (xt ^ 3) / 6 + (xt ^ 5) * 3 / 40 + (xt ^ 7) * 5 / 112 + (xt ^ 9) * 35 / 1152 + (xt ^ 11) * 63 / 2816 + (xt ^
13) * 231 / 13312
End Function
```

##### Function ASINd(xt As Single) As Single 'calcul avec angle en degré

```
ASINd = xt + (xt ^ 3) / 6 + (xt ^ 5) * 3 / 40 + (xt ^ 7) * 5 / 112 + (xt ^ 9) * 35 / 1152 + (xt ^ 11) * 63 / 2816 + (xt ^
13) * 231 / 13312
ASINd = ASINd * 45 / Atn(1)
End Function
```

##### Function DEGenRD(xt As Single) As Single ''donne angle converti en radian

```
DEGenRD = xt * Atn(1) / 45
End Function
```

##### Sub PLANOUIEvv2(z1 As String, z2 As String, z3 As String, vv As Single, deci As Integer)

```
sk.Seek "=", z1, z2, z3: If sk.NoMatch Then Stop
vv = Int(vv / 10 ^ deci + 0.5) * 10 ^ deci
sk.Edit
  sk("mm") = IIf(vv <> sk("vv2"), 1, 0) 'pour marquer un chgt de valeur donc mise en rouge
   sk("vv2") = vv
sk.Update
End Sub
```

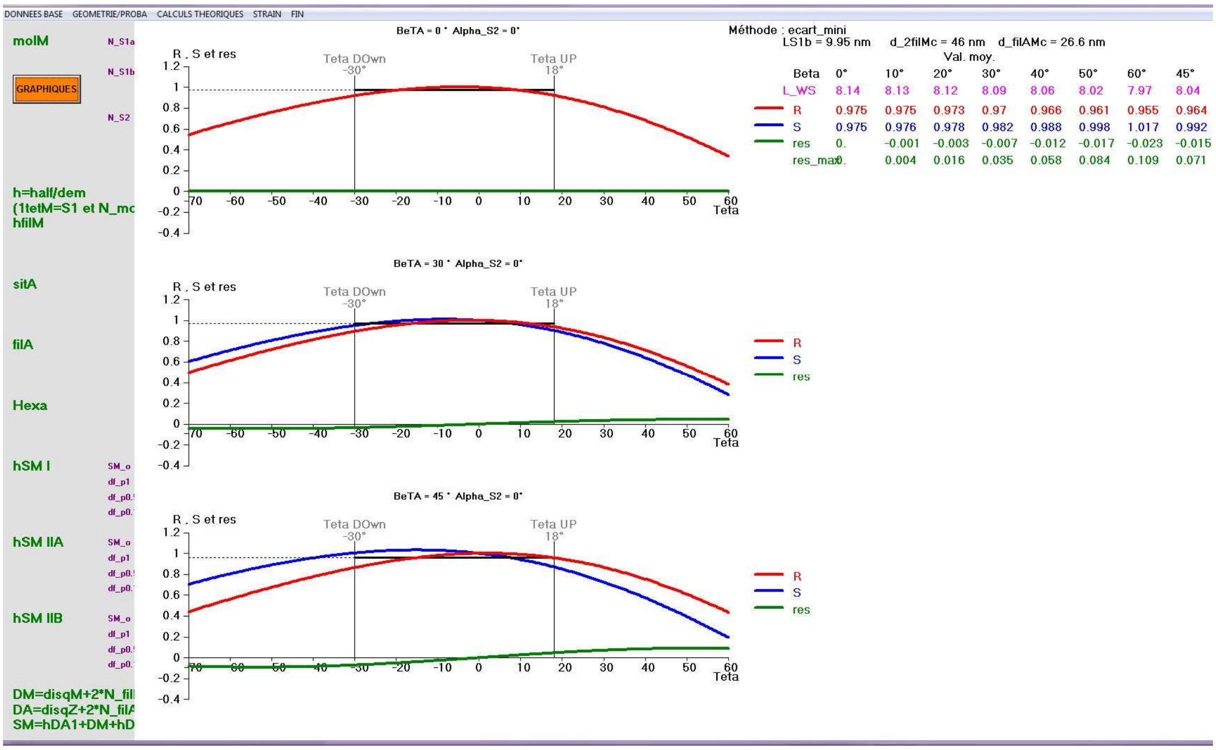


**Fig CP2.2. Screenshot after starting the "Sub AA_VIT_raccst_tetM() for plotting relationship between R, S and res as a function of θ.**

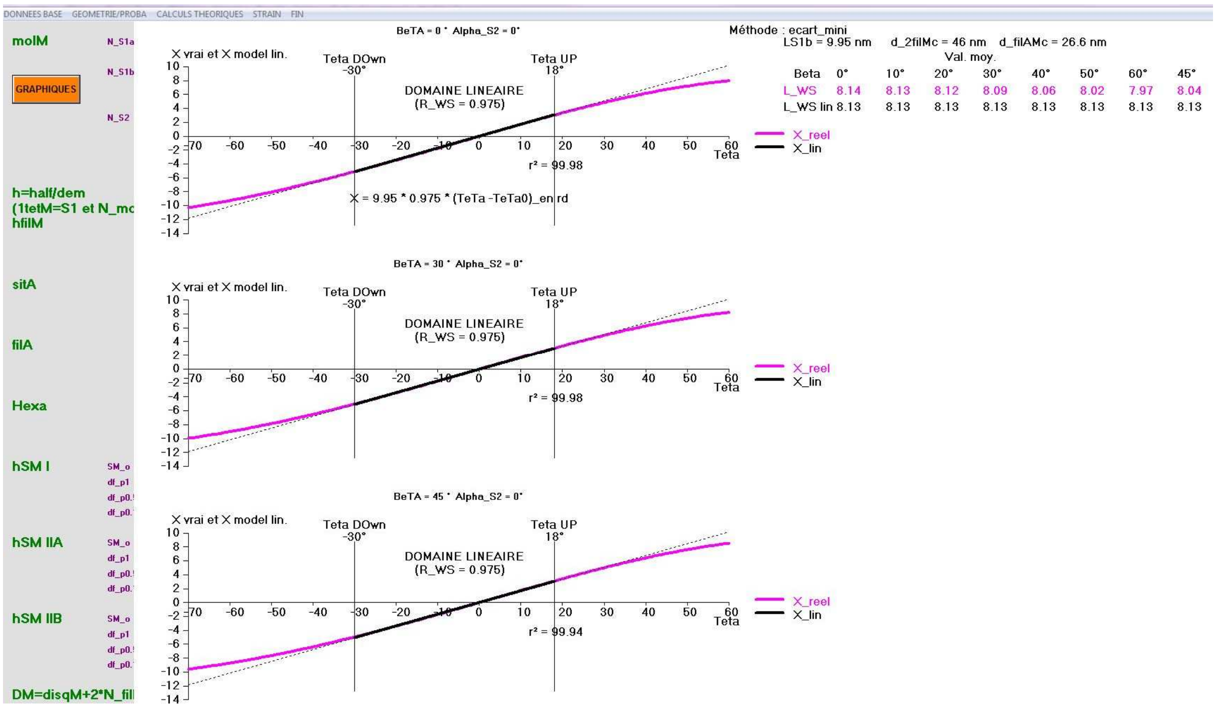


**Fig CP2.3. Screenshot after starting the "Sub AA_VIT_raccst_tetM() for plotting relationship between $\delta X_{eq}$ and $\delta X_{lin}$ as a function of θ.**

#### Sub VIT_raccsst_tetM()

```
'pi = Atn(1) * 4
L_WS = 11 'nm
xx = 30 + 30 'ds un hsR
xn = -40 - 30
'petite verif
If xn > xx Then Stop
xt = 10
'Teta0 = 8
vx = "Teta"

sk.Index = "kikekoi" ' entete, koi,lb
'sk.Seek "=", "molM", "S2", "BeTA": If sk.NoMatch Then Stop
Beta1 = 0 'sk("vv1")
'If Beta1 <> 0 And Beta1 <> 10 And Beta1 <> 20 And Beta1 <> 30 And Beta1 <> 40 And Beta1 <> 50 Then
Stop

sk.Seek "=", "molM", "S2", "Alpha": If sk.NoMatch Then Stop
ALPHA_S2 = sk("vv1") ' 0 'pour F = 1 à 3
ALPHA_S2rd = DEGenRD(ALPHA_S2)

sk.Seek "=", "molM", "S1b", "dTeTA": If sk.NoMatch Then Stop
dTeta_MAX = sk("vv1")

sk.Seek "=", "molM", "S1a", "Y": If sk.NoMatch Then Stop
YS1a = sk("vv2") 'S1a_Y=4 nm
sk.Seek "=", "molM", "S1b", "L": If sk.NoMatch Then Stop
LS1b = sk("vv1")
sk.Seek "=", "molM", "S2", "L": If sk.NoMatch Then Stop
LS2 = sk("vv1")
sk.Seek "=", "Hexa", "AM", "dY": If sk.NoMatch Then Stop
dY_PQ = sk("vv2") - YS1a 'd'apres donénes stockées "vrai dY_PQ" = 15 nm (entre 11 et 21 nm)


sk.Seek "=", "Hexa", "2filM", "d"
d_2filMc = sk("vv1") 'd_2filM(centres)
d_filAMc = d_2filMc * Sqr(3) / 3  'distance filA/filM (centres)

'sk.Seek "=", "Hexa", "filA/filM", "d": d_filAMc = sk("vv2")
sk.Seek "=", "hfilM", "hfilM", "R": r_filM = sk("vv1") 'R_filM
sk.Seek "=", "filA", "filA", "R": r_filA = sk("vv1") 'R_filA
sk.Seek "=", "molM", "S2", "pas": If sk.NoMatch Then Stop
xM = sk("vv2")

'********
'hentete = 40: hpage = 12
 X0 = -10 '-10IIf(mmm = 1, -15, -25)  'decalge vers la droite pour misen page impression
 'Y0 = -20
 larg = 200 ' '115IIf(mmm = 1, 140, 130) ' 140 'IIf(CYCIMP!Chi(3).Value = 0, 140, 170)
 'haut = 140
 'hentete = 7: hpage = 20
'********************************
BetaL = 45 '47'° limite WS

Xd = 0
XS1a = -2 ' a priroi on s'en fout

Set TeTa_X = dbM.OpenRecordset("TETAfonctionX", dbOpenTable): TeTa_X.Index = "PK"
```

```
'on efface tout ds table ang
dbM.Execute "delete * from TETAfonctionX"

'**** 3 chois a faire
'choix1 : quel model pour detremeiner tetaup et teta Down
   'model = "max_R" rejetté
   'model = "pente_0" 'retenu
   model = "ecart_mini" 'presque pareil que pente_0
'choix 2 : quel courbes ?
   'tx4 = "R_S_res"
   tx4 = "X_vrai et X_model"
   If tx4 = "R_S_res" Then
     yn = -0.4: yx = 1.2: yt = 0.2
     vy = "R , S et res"
   Else
      yn = -14: yx = 10: yt = 2
      vy = "X vrai et X model lin."
   End If
'choix 3
   tx3 = "0" ' beta =0°30°45°
   'tx3 = "60" 'beta =40°50°60°

'N_F = 3 'ou 6
For F = 1 To 3 'N_F ' 3 ou graphes

 Select Case F
  Case 1
    impscale X0, Y0 + haut + hentete, X0 + larg, Y0 - 2 * haut - hpage  'F=1à3
    VIT_raccsst_tetM_TETAfonctionX 'beta1=0 par deafut
    Beta1 = IIf(tx3 = "0", 0, 10) '30 '20
   Case 2
     impscale X0, Y0 + 2 * haut + hentete, X0 + larg, Y0 - haut - hpage  'F=1 à 3
      Beta1 = IIf(tx3 = "0", 30, 20) ' Beta1 = 20 '30 '20
   Case 3
     impscale X0, Y0 + 3 * haut + hentete, X0 + larg, Y0 - hpage 'F=1 à 3
      Beta1 = IIf(tx3 = "0", 45, 60) 'Beta1 = 40 '40
  End Select

   'titre à GAUCHE
    zY1 = "BeTA = " + Format(Beta1) + " ° Alpha_ S2 = " + Format(ALPHA_S2) + "°"
    O.ForeColor = QBColor(0): O.FontBold = -1: O.FontSize = 11 'IIf(mmm = 1, 10, 9)
    O.CurrentX = 50 - O.TextWidth(zY1) / 2: O.CurrentY = IIf(mmm = 1, 125, 127)
    O.Print zY1

   impenvi
   If tx4 = "R_S_res" Then
      VIT_raccsst_tetMpt "S", 9
       VIT_raccsst_tetMpt "R", 12
       VIT_raccsst_tetMpt "res", 2
   Else
     VIT_raccsst_tetMpt "X_reel", 13
     VIT_raccsst_tetMpt "X_lin", 0
   End If

Next 'f


End Sub
```

#### Sub VIT_raccsst_tetMpt(iX As String, zcoul As Integer)

```
Dim RouS As Single
'RouS = 0.94
'ran temporaria teta=14.33 exp(0.1916 X) --> X = 5.22 log(teta/14.33)   valable pour beta = 0°to 60°
'rana exulenta  teta=14.64 exp(0.2264 X) --> X = 4.417 log(teta/14.64)  valable pour beta = 0°to 60°
'sX = IIf(LS1b = 9, 14.64, 14.33) '14.64 '14.33=0.0698
'sY = IIf(LS1b = 9, 4.42, 5.22) '4.42 '1/01816=5.22
'rappels: TeTA=DEGenRD(xn)  et fi=calculFIrd(xn)

'*********integration/somme*****
'EPv = EPv + (pEPdt + avpEPdt) / (2 * Hzimadb)
'***************
'TeTA en X varie de xn=TeTA1enDEG à xx=Teta3enDEG

With O

TeTa_X.Seek "=", -360: If TeTa_X.NoMatch Then Stop
If ALPHA_S2 = 0 Then
    If tx4 = "R_S_res" Then
        RouS = TeTa_X("R_" + Format(Beta1))
    Else
        RouS = TeTa_X("R_0") 'opn prend toujours le m^me
    End If
Else
   If LS1b = 9.95 Then
      RouS = 0.944
   Else
      Stop ' a faire ou verifier
   End If
End If
If iX = "R" Then
   ymi = (RouS - yn) * 100 / (yx - yn)
   'traces teta 0tetaup et tetadown

   .ForeColor = QBColor(8)
   For P = 1 To 2
     xav = Choose(P, TetaDO, TetaUP)
     xav = (xav - xn) * 100 / (xx - xn)
     yav = (0 - yn) * 100 / (yx - yn)
     'trait vertical
     .DrawStyle = 0: .DrawWidth = 1
     O.Line (xav, yav)-(xav, 95), QBColor(0)
     txV = Choose(P, "Teta DOwn", "Teta UP")
     .CurrentX = xav - .TextWidth(txV) / 2
     .CurrentY = 109
     O.Print txV
     txV = Format(Choose(P, TetaDO, TetaUP)) + "º"
     .CurrentX = xav - .TextWidth(txV) / 2
     .CurrentY = 102
     O.Print txV
     'trait horizontal  R_mpoy=0.94
            If P = 1 Then
             .DrawStyle = 2
             xmi = 0
             xap = xav
            Else
             .DrawWidth = 3
             xmi = xap
```

```
            xap = xav
          End If
           O.Line (xmi, ymi)-(xap, ymi), QBColor(0)
   Next

  ElseIf iX = "X_reel" Then
     'calul X0_E
         'calul du vrai teta0
         'Teta2 = Int(TetaUP - dTeta_MAX * 7 / 10)
         'Teta0 = (TetaUP + Teta2) / 2
         'ou tout betement 0
         Teta0 = 0
         TeTa_X.Seek "=", Teta0
         X0_E = TeTa_X("X_" + Format(Beta1))
      .DrawWidth = 1
     For P = 1 To 2
        xav = Choose(P, TetaDO, TetaUP)
        xav = (xav - xn) * 100 / (xx - xn)
        'yav = (0 - yn) * 100 / (yx - yn)
       'trait vertical
        O.Line (xav, 5)-(xav, 95), QBColor(0)
        txV = Choose(P, "Teta DOwn", "Teta UP")
        .CurrentX = xav - .TextWidth(txV) / 2
        .CurrentY = 109
        O.Print txV
        txV = Format(Choose(P, TetaDO, TetaUP)) + "°"
        .CurrentX = xav - .TextWidth(txV) / 2
        .CurrentY = 102
         O.Print txV
      Next
End If

'tracé demandé
e_tt = 4 '1'6
.DrawWidth = e_tt

If iX = "X_lin" Then
   .DrawWidth = 1: .DrawStyle = 2
    xav = 0
    yav = LS1b * RouS * DEGenRD(xn - Teta0): yav = (yav - yn) * 100 / (yx - yn)
    xap = 100
    yap = LS1b * RouS * DEGenRD(xx - Teta0): yap = (yap - yn) * 100 / (yx - yn)
    O.Line (xav, yav)-(xap, yap), QBColor(zcoul)

   .DrawWidth = e_tt: .DrawStyle = 0
    xav = (TetaDO - xn) * 100 / (xx - xn)
    yav = LS1b * RouS * DEGenRD(TetaDO - Teta0): yav = (yav - yn) * 100 / (yx - yn)
    xap = (TetaUP - xn) * 100 / (xx - xn)
    yap = LS1b * RouS * DEGenRD(TetaUP - Teta0): yap = (yap - yn) * 100 / (yx - yn)
    O.Line (xav, yav)-(xap, yap), QBColor(zcoul)

ElseIf iX = "X_reel" Then
     tx1 = "X_" + Format(Beta1)
      'poit de départ
    TeTa_X.Seek "=", xn
    .CurrentX = 0 '(Pmin - xn) * 100 / (xx - xn)
     ymi = TeTa_X(tx1) - X0_E
    .CurrentY = (ymi - yn) * 100 / (yx - yn)
```

```
    sN = 0: sX = 0: sXX = 0: sY = 0: sYY = 0:  sXY = 0
    TeTa_X.MoveNext
    Do Until TeTa_X.EOF
        xap = TeTa_X("Teta")
        yap = TeTa_X(tx1) - X0_E
        If xap >= TetaDO And xap <= TetaUP Then
            sN = sN + 1
            sX = sX + yap: sXX = sXX + yap ^ 2
            ymi = LS1b * RouS * DEGenRD(xap - Teta0)
            sY = sY + ymi: sYY = sYY + ymi ^ 2
            sXY = sXY + yap * ymi
        End If
        xap = (xap - xn) * 100 / (xx - xn)
        yap = (yap - yn) * 100 / (yx - yn)
        O.Line -(xap, yap), QBColor(zcoul)
        TeTa_X.MoveNext
    Loop

     'r  regression classique
     'r_regL = (sXY - sX * sY / sN) / Sqr((sXX - sX * sX / sN) * (sYY - sY * sY / sN))

     'calcul ou la droit passe par zer0
     p_regL = sXY / sXX
     r_regL = 1 - (sYY - 2 * p_regL * sXY + sXX * p_regL ^ 2) / (sYY - sY * sY / sN)

     .CurrentX = 63: .CurrentY = 45
     O.Print "r² = " + Format(100 * r_regL ^ 2, "###.##")
     'O.Print "r2 = " + Format(r_regL ^ 2)

Else
    tx1 = iX + "_" + Format(Beta1)
    'poit de départ
    TeTa_X.Seek "=", xn
    .CurrentX = 0 '(Pmin - xn) * 100 / (xx - xn)
    .CurrentY = (TeTa_X(tx1) - yn) * 100 / (yx - yn)
    TeTa_X.MoveNext
    Do Until TeTa_X.EOF
        xap = (TeTa_X("Teta") - xn) * 100 / (xx - xn)
        yap = (TeTa_X(tx1) - yn) * 100 / (yx - yn)
        O.Line -(xap, yap), QBColor(zcoul)
        TeTa_X.MoveNext
    Loop
End If

'************** LEGENDES
  xav = 105
  Select Case iX
    Case "R": yav = 75 'Choose(Ix, 65, 75, 40, 85, 70, 60, 50)
    Case "S": yav = 65
    Case "res": yav = 55
    Case "X_reel": yav = 60
    Case "X_lin": yav = 52
    Case Else: Stop
  End Select
  O.Line (xav, yav)-(xav + 5, yav), QBColor(zcoul)
  .ForeColor = QBColor(zcoul): .FontBold = -1
  .CurrentX = xav + 7
  .CurrentY = yav + 3
```

```
  O.Print iX

'*** affichage Rmoy et Smoy
  xav = 120
  TeTa_X.Seek "=", -360
  If F = 1 Then
     If iX = "R" Or iX = "X_lin" Then
       .ForeColor = QBColor(0)
       .CurrentX = 100: .CurrentY = 126: O.Print "Méthode : " + model
       .CurrentX = 110: .CurrentY = 119: O.Print "LS1b = " + Format(LS1b) + " nm"
       .CurrentX = 130: .CurrentY = 119: O.Print "d_2filMc = " + Format(d_2filMc) + " nm"
       .CurrentX = 150: .CurrentY = 119: O.Print "d_filAMc = " + Format(d_filAMc, "##.#") + " nm"
       .CurrentX = 140: .CurrentY = 110: O.Print "Val. moy."
       'totue les vls mot pour beta dee 0+ à 60°
       .CurrentX = 112: .CurrentY = 100: O.Print "Beta"
       .ForeColor = QBColor(13)
       .CurrentX = 110: .CurrentY = 90: O.Print "L_WS"

       For zB = 0 To 7 'Beta= 0°à 60°
         'BETA
         z = IIf(zB < 7, zB * 10, BetaL) '  10°20 °... 60°45°
         'zp = zB * 9 'position
          .ForeColor = QBColor(0)
         .CurrentX = xav + zB * 9
         .CurrentY = 100
         O.Print Format(z) + "°"
         'L_WS vrai
         .ForeColor = QBColor(13)
         .CurrentX = xav + zB * 9
         .CurrentY = 90
          O.Print Format(TeTa_X("X_" + Format(z))) '+ "nm"
       Next
    End If

    If tx4 = "R_S_res" Then
       If iX = "res" Then
         .CurrentX = 112
         .CurrentY = yav - 7
        .ForeColor = QBColor(zcoul)
         O.Print "res_max"
       End If

      For zB = 0 To 7 'Beta= 0 à 50°
        .ForeColor = QBColor(0)
         z = IIf(zB < 7, zB * 10, BetaL) '  10°20 °... 60°45°
         'zp = zB * 9 'position
        txV = iX + "_" + Format(z)
        .CurrentX = xav + zB * 9
        .CurrentY = yav + 3
        .ForeColor = QBColor(zcoul)
        O.Print Format(TeTa_X(txV), "0.###")

        If iX = "res" Then 'res_max
          .CurrentX = xav + zB * 9
          .CurrentY = yav - 7
           O.Print Format(TeTa_X(txV + "_Max"), "0.###")
        End If
      Next
```

```
   End If
 End If

 If iX = "X_lin" Then
      'ymi = LS1b * RouS * DEGenRD(P - Teta0)
        .ForeColor = QBColor(zcoul)
        .CurrentX = 40: .CurrentY = 90
         O.Print "DOMAINE LINEAIRE"
        .CurrentX = 42: .CurrentY = 82
         O.Print "(R_WS = " + Format(RouS, "0.###") + ")"

        If F = 1 Then

         .CurrentX = 30: .CurrentY = 25
         O.Print "X = " + Format(LS1b) + " * " + Format(RouS, "0.###") + " * (TeTa -TeTa0)_en rd"

        .CurrentX = 110: .CurrentY = 80: O.Print "L_WS lin"
        For zB = 0 To 7 'Beta= 0°à 60°
          ' z = IIf(zB < 7, zB * 10, betal)
          .CurrentX = xav + zB * 9
          .CurrentY = 80
           O.Print Format(LS1b * RouS * DEGenRD(dTeta_MAX), "##.##") ' + "nm"
        Next
        End If

 End If

End With
End Sub
```

#### Sub impscale(X1 As Integer, y1 As Integer, X2 As Integer, Y2 As Integer)

```
           O.ScaleLeft = X1
           O.ScaleTop = y1
           O.ScaleWidth = X2 - X1
           O.ScaleHeight = Y2 - y1
End Sub
```
