## Supplementary Chapter. Forces and energies involved in the working stroke of a myosin II head. for "Mechanical model of muscle contraction 2. Kinematic and dynamic aspects of a myosin II head during the working stroke"

### S2.C Supplementary chapter of Paper 2

### Forces and energies related to a myosin head in working stroke

A myosin II head is modelled by 3 segments, the motor or catalytic domain (S1a), the lever (S1b) and the rod (S2). A head in working stroke (WS) is characterized by: (a) the rigidity of the 3 segments; (b) the strong binding between S1a and the actin molecule; (c) the motor moment ($\mathcal{M}_B$) exerted on S1b, a moment which induces a traction of the myosin filament via S2, leading *de facto* to a shortening of the half-sarcomere (hs); (d) the displacement of S1b in a fixed plane, the orientation of S1b in this plane being defined with the angle θ (Hypothesis n° 4 of the model formulated in Paper 2).

#### C.1 Units at the scale of a myosin II head

The weight unit is kiloDalton (kDa) or yoctoKilogram (yKg):

$$1\ \text{kDa} = 1.66\ 10^{-24}\ \text{Kg} = 1.66\ \text{yKg}$$

The unit of length is the nanometer (nm):

$$1\ \text{nm} = 10^{-9}\ \text{m}$$

The unit of velocity is the nanometer per millisecond ($\text{nm}\cdot\text{ms}^{-1}$):

$$1\ \text{nm}\cdot\text{ms}^{-1} = 10^{-3}\ \text{mm}\cdot\text{s}^{-1} = 10^{-6}\ \text{m}\cdot\text{s}^{-1}$$

The unit of force is the picoNewton (pN):

$$1\ \text{pN} = 10^{-12}\ \text{N}$$

The unit of energy is the zeptoJoule (zJ):

$$1\ \text{zJ} = 1\ \text{pN} \cdot 1\ \text{nm} = 10^{-21}\ \text{J}$$

#### C.2 Brownian displacements and thermal shocks

A large molecule about ten nanometers in size is placed in a liquid. It undergoes the multiple shocks of the surrounding small molecules, components of the liquid. Brownian agitation corresponds to the fluctuation in the number of microscopic states of the large molecule and therefore to the fluctuation in its entropy. In 1910, A. Einstein calculated that this fluctuation was equal to $k_B/2$ where $k_B$ is the Boltzmann constant. Each shock creates an energy variation ($\delta E_{th}$) equal to:

$$\delta E_{th} = \frac{k_B \cdot \Gamma_{amb}}{2} \qquad \text{(C1)}$$

where $k_B = 1.38\cdot10^{-23}\ \text{J}\cdot°\text{K}^{-1}$ ; $\Gamma_{amb}$ is the temperature of the ambient liquid expressed in °K.

In muscle physiology, experimental temperatures verify:

$$0\ °\text{C} \leq \Gamma_{amb} \leq 35\ °\text{C}$$

This implies:

$$3.77\ \text{zJ} \leq (k_B \cdot \Gamma_{amb}) \leq 4.25\ \text{zJ}$$

Using (C1) and previous inequalities, the average value of the energy fluctuation during a thermal shock is deduced:

$$\delta E_{th} \approx 2 \pm 0.12 \text{ zJ} \quad \text{(C2)}$$

**C.3 Hydrolysis of an ATP molecule and maximum tensile force of a myosin II head**

According to the thermodynamic tables, the free enthalpy variation of ATP hydrolysis under physiological conditions is between -50 and -40 kJ per mole of ATP, either for an ATP molecule by dividing by $6.023 \cdot^{1023}$, the Avogadro number, a value between -83 and -66 zJ. The chemical energy resulting from the hydrolysis of an ATP molecule is partially converted into mechanical energy stored as elastic potential energy ($EP_{S1}$) in the β-sheet element of the motor domain (S1a). By choosing a maximum output in the range of 40% to 60% [1], we obtain as a range of maximum values for $EP_{S1}$ :

$$(0.4 \text{ x } 66 \text{ zJ}) \leq EP_{S1,Max} \leq (0.6 \text{ x } 83 \text{ zJ})$$

So:

$$25 \text{ zJ} \leq EP_{S1,Max} \leq 50 \text{ zJ} \quad \text{(C3)}$$

According to (C2), $EP_{S1,Max}$ is 13 to 25 greater than $\delta E_{th}$.

The motor-moment ($\mathcal{M}_B$), a function of the θ angle, is derived from $EP_{S1}$:

$$\mathcal{M}_B(\theta) = -\frac{d}{d\theta}(EP_{S1}) \quad \text{(C4)}$$

After studying phase 1 of a length step, it is established that the motor-moment during the WS is a decreasing function of the angle θ. Some authors have suggested that the relationship is linear [2,3,4] We will follow them with hypothesis 6 presented in Paper 4 where the relationship between motor moment ($\mathcal{M}_B$) and θ is formulated:

$$\mathcal{M}_B(\theta) = \mathcal{M}_{up} \cdot \frac{(\theta - \theta_{down})}{\delta\theta_{Max}} \quad \text{(C5)}$$

where $M_{up}$ is the maximum moment corresponding to the angle $\theta_{up}$, i.e. the referent indicator of the maximum chemical-mechanical efficiency of the hydrolysis reaction of an ATP molecule previously studied; $\delta\theta_{Max}$ is the angular interval between $\theta_{up}$ and $\theta_{down}$; the values of these 2 angles being calculated respectively in (12a) and (12b) in Paper 2.

After integration of the relationship (C5), we obtain the expression of $EP_{S1}$:

$$EP_{S1}(\theta) = \mathcal{M}_{up} \cdot \frac{(\theta_{down} - \theta)^2}{2 \cdot \delta\theta_{Max}} \quad \text{(C6)}$$

Equation (16) of Paper 2 provides the relationship between the tensile force of a WS head and the motor-moment exerted on the lever. The maximum tensile force ($T_{S1,Max}$) is deduced from this:

$$T_{S1,Max} = \frac{\mathcal{M}_{up}}{L_{S1b} \cdot S_{WS}}$$

where $L_{S1b}$ is the lever length; $S_{WS}$ is a constant calculated in equations (13) and (14) of Paper 2.

Otherwise formulated with (C6):

$$T_{S1,Max} = \frac{2 \cdot EP_{S1,max}}{S_{WS} \cdot \delta\theta_{Max} \cdot L_{S1b}} \tag{C7}$$

where the constants $\delta\theta_{Max}$ and $S_{WS}$ are given equal, respectively, to 70° and 0.945 according to the equalities (8) and (13) of Paper 2.

With (C3) and (C7), we check that the value of $T_{S1,Max}$ varies according to that of $L_{S1b}$:

$$L_{S1b} = 9 \text{ nm} \quad \Rightarrow \quad 5 \text{ pN} \leq T_{S1,Max} \leq 10 \text{ pN} \tag{C8a}$$

$$L_{S1b} = 10 \text{ nm} \quad \Rightarrow \quad 4.5 \text{ pN} \leq T_{S1,Max} \leq 9 \text{ pN} \tag{C8b}$$

$$L_{S1b} = 11 \text{ nm} \quad \Rightarrow \quad 4 \text{ pN} \leq T_{S1,Max} \leq 8 \text{ pN} \tag{C8c}$$

These values are compatible with those of the literature [5,6].

From the above, we deduce the condition:

$$T_{S1,Max} \geq 4 \text{ pN} \tag{C9}$$

#### C.4 Strong Binding (SB)

When S1a is strongly attached to the actin molecule, the associated state is called *strong binding*, and adopts the acronym SB. Equation (4) of Paper 2 gives a relation of equality between the tangential components of the forces exerted on the actin filament (Afil) and on the myosin filament (Mfil), which means that the SB force ($F_{SB}$) between S1a and actin molecule also checks (C9):

$$F_{SB} \geq 4 \text{ pN} \tag{C10}$$

#### C.5 Weak Binding (WB)

When S1a is weakly attached to the actin molecule, the corresponding state is called *weak binding*, and takes the acronym WB. During the WB state, S1a undergoes the shocks of the molecules of the intracellular liquid and follows a Brownian trajectory in a volume circumscribed by the return force of the weak binding ($F_{WB}$) exerted by the actin molecule. The energy to break the WB ($\delta E_{WB}$) must be greater than or equal to $3 \cdot \delta E_{th}$:

$$\delta E_{WB} \geq 3 \cdot \delta E_{th} \tag{C11}$$

The factor 3 comes from the fact that shocks generate displacements of S1a and S1b according to the 3 axes of space and it is necessary to multiply by 3 with respect to the law of equipartition of energy.

When the WB is broken, there is a possibility of displacement of the head between two actin molecules adjacent and distant from $L_{Amol,}$ a distance equivalent to the diameter of actin molecule. The calculation of work as the product of force ($F_{WB}$) by displacement ($L_{Amol}$) gives according to (C11):

$$F_{WB} \geq \frac{3 \cdot \delta E_{th}}{L_{molA}} \quad \text{(C12)}$$

The expression of the work of a force equal to the product force by length is valid if the force is constant, which is not the case for an electrostatic attraction force that varies with distance. Inequality (C12) provides an average order of magnitude. With (C2) and $L_{Amol}$ = 5.5 nm (see Table 1 of accompanying Paper 3), it is verified:

$$F_{WB} \geq 1 \text{ pN} \quad \text{(C13)}$$

As by nature $F_{WB}$ must be less than $F_{SB}$, we deduce from (C10) and (C13):

$$1 \text{ pN} \leq F_{WB} \leq 4 \text{ pN} \quad \text{(C14)}$$

#### C.6 Acceleration quantities of gravitational origin

From data in the literature [7], it is possible to calculate an approximate weight for the different components of the muscle fiber, and to calculate for each of them the corresponding gravitational force.

The calculation of the force due to earth gravity acting on lever S1b ($P_{S1b}$) gives:

$$P_{S1b} = M_{S1b} \cdot g = 70 \cdot 1.66 \cdot 10^{-24} \cdot 9.81 \text{ Kg} \cdot \text{m} \cdot \text{s}^{-2} = 10^{-9} \text{ pN}$$

where $M_{S1b}$ is the mass of S1b equal to about 70 kDa; g is the acceleration due to gravity that is 9.81 $m \cdot s^{-2}$ on the earth's surface.

The force due to earth gravity acting on the actin filament ($P_{Afil}$) is equal to:

$$P_{Afil} = M_{Afil} \cdot g = 25 \cdot 10^{3} \cdot 1.66 \cdot 10^{-24} \cdot 9.81 \text{ Kg} \cdot \text{m} \cdot \text{s}^{-2} = 4\ 10^{-7} \text{ pN}$$

where $M_{Afil}$ is the mass of an Afil equal to about $25 \cdot 10^3$ kDa.

The value of force due to earth gravity exerted on a hs ($P_{hs}$) is:

$$P_{hs} = M_{hs} \cdot g = 10^{7} \cdot 1.66 \cdot 10^{-24} \cdot 9.81 \text{ Kg} \cdot \text{m} \cdot \text{s}^{-2} = 1.6\ 10^{-4} \text{ pN}$$

where $M_{hs}$ is the mass of a hs whose maximum value is in the order of $10^7$ kDa.

For these 3 examples, the calculated forces are negligible in relation to internal forces of elastic or electrostatic origin whose values are in the order of pN. At the nanoscale, the influences of the potential energy of gravitational origin and its variations will be considered as zero, regardless of the position of the isolated fiber in relation to the earth's gravity field during experiments. A practical consequence is that the force exerted by a muscle does not depend on its position in space.

#### C. 7 Acceleration quantities of inertial origin

The relative maximum shortening velocity per half-sarcomere ($u_{Max}$) is calculated during phase 1 of a 10 nm length step during 100 µs as:

$$u_{Max} = 100\ nm \cdot ms^{-1} = 10^{-4}\ m \cdot s^{-1}$$

In a reference frame linked to any M-disk and assumed Galilean, the segment likely to undergo the greatest quantities of linear and angular acceleration is the lever S1b. When the relative velocity changes from 0 to $u_{Max}$, the maximum variation in the kinetic energy of the lever ($\Delta K_{\mathbf{S1b}}$) is equal to:

$$\Delta K_{S1b} = \frac{1}{2} \cdot M_{S1b} \cdot u_{Max}^{2} + \frac{1}{2} \cdot J_{S1b} \cdot \dot{\theta}_{Max}^{2}$$

where $J_{S1b}$ is the moment of inertia of a rod:

$$J_{S1b} = M_{S1b} \cdot \left( \frac{r_{S1b}^{2}}{4} + \frac{L_{S1b}^{2}}{12} \right) \approx 70 \cdot 1.66 \cdot 10^{-24} \cdot \left( \frac{1.5^{2}}{4} + \frac{10^{2}}{12} \right) \cdot 10^{-18} = 10^{-39}\ Kg \cdot m^{2}$$

and where $\dot{\theta}_{Max}$ is the maximum angular velocity of S1b, calculated as follows:

$$\dot{\theta}_{Max} \approx \frac{u_{Max}}{L_{S1b}} = 10^{4}\, rd \cdot s^{-1}$$

The result is:

$$\Delta K_{S1b} \approx 10^{-10}\, zJ$$

This is a negligible energy corresponding to acceleration forces of less than $10^{-11}$ pN.

**Conclusion:** At the nanoscale, the quantities of linear and angular accelerations of inertial origin are negligible.

#### C. 8 Archimedes' Forces

It is similarly verified that the forces due to Archimedes' thrust are negligible.

For example, the calculation of Archimedes' force relative to S1a ($F_{Arch,S1a}$) gives:

$$F_{Arch,S1a} = V_{S1a} \cdot \rho_{H2O} \cdot g$$

where $\rho_{water}$ is the density of water equal to $10^{3}\ Kg \cdot m^{-3}$

$V_{S1a}$ is the volume of S1a such that:

$$V_{S1a} = \pi \cdot r_{S1a}^{2} \cdot L_{S1a} = 3.14 \cdot (3\ nm)^{2} \cdot 5\ nm \approx 10^{-25}\ m^{3}$$

After calculation:

$$F_{Arch,S1a} = 10^{3} \cdot 10^{-25} \cdot 9.81\ Kg \cdot m \cdot s^{-2} = 10^{-9}\ pN$$

#### C.9 Viscous type friction forces

The molecules that make up a viscous medium exert a resistance force on any moving object. The viscous force is calculated according to the Stokes-Einstein formula. It implies for segment S1a:

$$F_{Visc,S1a} = -(6 \cdot \pi \cdot \eta \cdot r_{S1a}) \cdot V_{S1}$$

where $\eta$ is the viscosity coefficient of the physiological liquid in which S1a moves; $r_{S1a}$ is the radius of the contact volume with the liquid displaced during the movement of S1a; $V_{S1}$ is the absolute velocity of the subfragment S1 in the laboratory reference frame, identical velocity for S1a and S1b.

With $\eta_{3°C} = 10^{-3}$ Kg.m$^{-1}$.s$^{-1}$, $r_{S1a}$ = 2 nm, $V_{S1}$= 2.5 mm.s$^{-1}$, the following calculation is performed:

$$F_{Visc,S1a} = -(6 \cdot 3.14 \cdot 10^{-3} \cdot 2 \cdot 10^{-9}) \cdot 2.5 \cdot 10^{-3} \text{ N} \approx 0.1 \text{ pN}$$

The same applies to S1b:

$$F_{Visc,S1b} = -(6 \cdot \pi \cdot \eta \cdot r_{S1b}) \cdot V_{S1} \approx 0.1 \text{ pN}$$

We note that these two forces are of the same order of magnitude, that they are proportional to $\eta$ and decrease as the temperature increases, that they oppose the rotation movement of S1b in a half-sarcomere on the right and accompany it in a half-sarcomere on the left. Moreover, the calculation only concerns the most distal hs: the closer we get to the fixed end of the myofibril, the more these 2 viscous forces decrease.

It will be assumed that viscosity forces do not influence myosin heads, as already observed in Paper 1 with Fig 3 where the initial length of the sarcomere does not influence the maximum velocity of shortening.

#### C.10 Conclusion

At the nanoscale, the quantities of linear and angular accelerations, of gravitational or inertial origin, are negligible; the same applies to the Archimedes thrust and the viscosity forces acting on the myosin heads. The only actions present in the mechanical system proposed by this model are the inter-segmental linking forces and moments of the WS heads, with two exceptions:

1/ during the phase 1 of a length or force step, in lenghtening or shortening, the viscosity forces are applied significantly to the solids constituted by the Z-disks and the M-disks associated, respectively, with the inter-digited actin and myosin filaments, (see Supplement S4.J of accompanying Paper 4).

2/ during the phase 4 of a force step for the high shortening velocities (see Paper 1).

#### References of Supplement S2.C


1. **Barclay CJ, Woledge RC, Curtin NA (2010)** Inferring crossbridge properties from skeletal muscle energetics. Prog Biophys Mol Biol 102: 53-71.
2. **Ford LE, Huxley AF, Simmons RM (1981)** The relation between stiffness and filament overlap in stimulated frog muscle fibres. J Physiol 311: 219-249.
3. **Piazzesi G, Dolfi M, Brunello E, Fusi L, Reconditi M, et al. (2014)** The myofilament elasticity and its effect on kinetics of force generation by the myosin motor. Arch Biochem Biophys 552-553: 108-116.
4. **Piazzesi G, Lucii L, Lombardi V (2002)** The size and the speed of the working stroke of muscle myosin and its dependence on the force. J Physiol 545: 145-151.
5. **Piazzesi G, Reconditi M, Linari M, Lucii L, Bianco P, et al. (2007)** Skeletal muscle performance determined by modulation of number of myosin motors rather than motor force or stroke size. Cell 131: 784-795.
6. **Tsaturyan AK, Bershitsky SY, Koubassova NA, Fernandez M, Narayanan T, et al. (2011)** The fraction of myosin motors that participate in isometric contraction of rabbit muscle fibers at near-physiological temperature. Biophys J 101: 404-410.
7. **Poortmans JR, Boisseau N (2002)** Biochimie des activités physiques: De Boeck Université.
